# Hydrogen Sulfide Coordinates Glucose Metabolism Switch through Destabilizing Tetrameric Pyruvate Kinase M2

**DOI:** 10.1101/2023.09.02.555034

**Authors:** Rong-Hsuan Wang, Pin-Ru Chen, Yue-Ting Chen, Yi-Chang Chen, Yu-Hsin Chu, Chia-Chen Chien, Po-Chen Chien, Shao-Yun Lo, Zhong-Liang Wang, Min-Chen Tsou, Ssu-Yu Chen, Guang-Shen Chiu, Wen-Ling Chen, Yi-Hsuan Wu, Lily Hui-Ching Wang, Wen-Ching Wang, Shu-Yi Lin, Hsing-Jien Kung, Lu-Hai Wang, Hui-Chun Cheng, Kai-Ti Lin

## Abstract

Cancer cells reprogram their glucose metabolic pathway from oxidative phosphorylation toward aerobic glycolysis. Pyruvate kinase M2 (PKM2), which converts phosphoenolpyruvate (PEP) to pyruvate, is considered the rate-limiting enzyme involved in cancer glucose metabolism. By reducing PKM2 enzyme activity, cancer cells attain a greater fraction of glycolytic metabolites for macromolecule synthesis needed for rapid proliferation. Here we demonstrate that hydrogen sulfide (H_2_S) destabilizes PKM2 tetramer into dimer/monomer, leading to reduced PKM2 enzyme activity and an increase in the activation of nuclear transcriptional genes mediated by dimeric PKM2. Proteomic profiling of endogenous PKM2 reveals the occurrence of sulfhydration at cysteines, notably at cysteine 326. Blocking PKM2 sulfhydration at cysteine 326 through amino acid mutation stabilizes PKM2 tetramer and crystal structure further indicating that the tetramer organization of PKM2^C326S^ is different from the currently known T or R states, revealing PKM2^C326S^ as a newly identified form. The presence of a PKM2^C326S^ mutant in cancer cells effectively rewires glucose metabolism to mitochondrial respiration, resulting in the significant inhibition of tumor growth. Collectively, PKM2 sulfhydration by H_2_S serves as a glucose metabolic rewiring mechanism in promoting tumorigenesis, and inhibition of PKM2 sulfhydration may be applied as a new therapeutic approach targeting cancer metabolism.

**One-Sentence Summary:** H_2_S rewires glucose metabolism by destabilizing PKM2 tetramerization majorly through sulfhydration at cysteine 326

**Highlights:**

1. H_2_S enhances PKM2 dissociation from tetramer to dimer to facilitate dimeric PKM2 nuclear translocation.
2. H_2_S modifies PKM2 sulfhydration, notably at cysteine 326.
3. The crystal structure reveals PKM2^C326S^ as a unique tetramer conformation.
4. Blockage of PKM2 sulfhydration at C326 rewires cancer glucose metabolism and significantly inhibits tumor growth.

## Introduction

Cancer metabolism refers to the reprogramming of cellular metabolic pathways in cancer cells to acquire the necessary nutrients to build new biomass. These metabolic rewiring activities in cancer cells are recognized as a general hallmark of cancer ^1^. Among the altered metabolic pathways, Warburg effect ^2^, a preferential dependence on aerobic glycolysis, is the classical example of cancer-specific metabolism. Although glycolysis is less efficient than oxidative phosphorylation regarding adenosine triphosphate (ATP) production, glycolysis provides metabolic intermediates required for the biosynthesis of nucleotides and amino acids, which cancer cells depend on for rapid cell proliferation under insufficient nutrient supply ^3^. The observation of this differential metabolic preference in cancer cells unveils a therapeutic potential by targeting this specific metabolic adaptation utilized by cancer cells. Indeed, drugs intended to block aerobic glycolysis in cancer cells are currently under evaluation, including the inhibitors against lactate dehydrogenase ^4^ and monocarboxylate transporters ^5^. Further studies of drugs targeting those tumor-specific metabolic pathways are highly warranted.

Pyruvate kinase (PK) regulates the final rate-limiting step of glycolysis by catalyzing the phosphoryl transfer from phosphoenolpyruvate (PEP) to adenosine diphosphate (ADP) to produce pyruvate and ATP ^6^. In mammals, two genes encode four mammalian PK isoforms (PKM1, PKM2, PKR, PKL). PKM2 is highly expressed in proliferating cells, including all types of cancer cells and tumors tested so far ^7^. Despite the sequence similarity, PKM1 and PKM2 have different catalytic properties. PKM1 is constitutively organized as a tetramer with high catalytic activity. In contrast, PKM2 maintains low activities in cancer cells through binding to various ligands or posttranslational modifications (PTMs) ^8^, including phosphorylation ^9^, acetylation ^10^, oxidation ^11^, SUMOylation ^12^, O-GlcNAcylation ^13^, and methylation ^14^. These modifications of PKM2 downregulate its enzymatic activity, thus redirecting the use of glucose from energy production to biomass synthesis for fast-proliferating tumor cells. In addition, the dimeric PKM2 also presents non-canonical functions in different cellular localization, such as nuclei, mitochondria, and exosomes, to support tumor growth and distant metastasis ^15^.

Hydrogen sulfide (H_2_S), a colorless, flammable, water-soluble gas, is an endogenously produced gasotransmitter that acts as a critical mediator in multiple physiological processes ^16^, including vasodilation ^17–19^, aging ^20^, and inflammation ^21^. Aberrant up-regulation of H_2_S synthesizing enzymes, cystathionine β-synthase (CBS), cystathionine γ-lyase (CTH), and 3-mercaptopyruvate sulfurtransferase (3-MST), is frequently observed in multiple cancer types ^22^. Endogenous H_2_S promotes tumor growth through anti-apoptosis, DNA repair, tumor growth, cancer metabolism, metastasis, and angiogenesis ^22^, mainly through the formation of a persulfide (-SSH) bond on Cysteine, which is called protein sulfhydration ^23^. Currently, the influences of this novel PTM remain largely unexplored, more investigations are expected to reveal how H_2_S impacts cancer progression.

PKM2 activity can be inhibited by different amino acids ^8^, in particular, L-cysteine ^24^. Since H_2_S is mainly produced from L-cysteine ^25^, we started to investigate whether PKM2 activity could be modulated by H_2_S. Here, we show that cancer cells utilize H_2_S to destabilize PKM2 tetramers into dimers or monomers with low PK activity. We further determined the sulfhydration sites on PKM2 by mass spectrometry. A single amino acid mutation on cysteine 326 stabilizes PKM2 tetramer with high PK activity, resulting in the rewiring of the glucose metabolism toward mitochondrial respiration, leading to increased cytokinesis failure, as well as reduced cell proliferation and tumor growth in the mouse xenograft model. Our data support the hypothesis that the sulfhydration is a key post-translational modification to modulate PKM2 activity. Cysteine 326 on PKM2 may serve as a therapeutic target for future anti-cancer drug development.

## Results

### The sulfhydration of PKM2 through H_2_S promotes the dissociation of the active tetramer PKM2 to less active monomer/dimer PKM2

To investigate the role of H_2_S on PKM2 activity through protein sulfhydration. We first confirmed that treatment with NaHS, the H_2_S donor, induced sulfhydration of PKM2 by using a modified biotin switch assay to pull down sulfhydrated proteins in breast cancer MDA-MB-231 cell lysates (Fig. 1A) and prostate cancer PC3 cell lysates (Fig. S1A). This modification was reversed after treatment of dithiothreitol (DTT), the reagent used to cleave the disulfide bonds (Fig. 1A). Meanwhile, the activities of PKM2 were reduced in the presence of NaHS in both recombinant proteins (Fig. 1B) and in cancer cells (Fig. S1B), and this inhibition was reversed by DTT treatment (Fig. 1B), suggesting disrupting sulfhydration by DTT can restore PK activity of PKM2. Next, to determine whether H_2_S-mediated sulfhydration of PKM2 is involved in the oligomerization of PKM2, we performed glycerol gradient ultracentrifugation on purified recombinant PKM2 to differentiate the tetramer and monomer/dimer forms of PKM2 (Fig. 1C). The majority of the PKM2 in solution was a mixture in both monomer/dimer form and tetramer form, while exposure of PKM2 to its endogenous activator, fructose 1,6-bisphosphate (FBP), resulted in a shift of monomer/dimer into the tetramer (Fig. 1C). Interestingly, treatment of NaHS resulted in the significant dissociation of FBP-induced tetramer (Fig. 1C), suggesting that allosteric regulation of PKM2 may be modulated by H_2_S-mediated sulfhydration. We further confirmed that FBP remained the same after NaHS treatment (Fig. S1C) to exclude the possibility that FBP may dissociate into fructose by NaHS. Studies have shown that the PKM2 in the dimer state can translocate into the nucleus to act as a transcriptional coactivator to stimulate the transcription of genes that are required for tumor growth ^26^. We next checked PKM2 nuclear translocation and expression of PKM2 response genes, including Cyclin D1 (CCND1), Hypoxia-inducible factor 1-α (HIF-1α), Lactate Dehydrogenase A (LDHA), Glucose transporter 1 (GLUT1), GLUT12, Hexokinase 2 (HK2), and Glutaminase-1 (GLS1). We found that the nuclear translocation of PKM2 was significantly enhanced by NaHS treatment (Fig. 1D), and PKM2-response genes were induced in the presence of NaHS (Fig. 1E). Moreover, treatment with the H_2_S inhibitor, aminooxyacetic acid (AOAA), demonstrated a reduction of PKM2 nuclear translocation (Fig. S1D), resulting in the inhibition of several PKM2-responsive genes (Fig. S1E). Notably, depletion of H_2_S supplement through AOAA significantly suppressed cell proliferation in MDA-MB-231 cells (Fig. 1F). Overall, our data strongly indicate that H_2_S-mediated sulfhydration modulates the dissociation of PKM2 tetramers into dimers/monomers, resulting in decreased PK activities and heightened transcriptional activities of PKM2 responsive genes.

**Figure 1.**
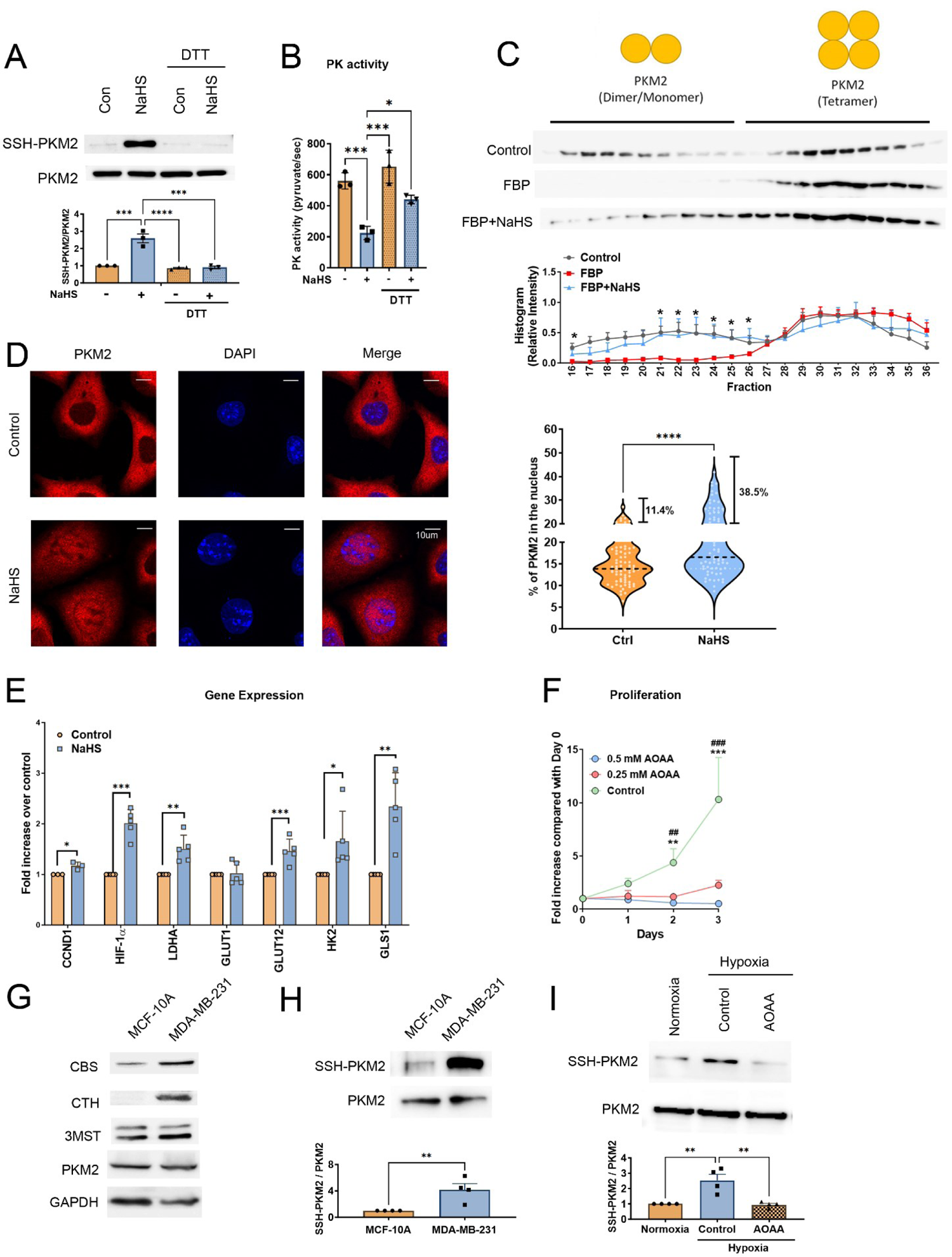
H_2_S destabilizes tetramerization of PKM2. (A) Upper: MDA-MB-231 cell lysates were treated with 100 μM NaHS for 30 min at 37°C and subjected to 1 mM DTT for 10 min. The biotin switch assay was then applied to precipitate sulfhydrated proteins. The biotin-labeled protein was analyzed by immunoblotting with anti-PKM2 antibody to detect sulfhydration of PKM2. Bottom: The SSH-labeled PKM2 was normalized with the level of total PKM2. Data are presented as the means ± SEM (n=3 biological replicates). One-way ANOVA followed by post hoc test was used for the statistical analysis (***p<0.001; ****p<0.0001). (B) The PK activity on recombinant PKM2 in the presence of 100 μM NaHS for 30 minutes on ice and subject to 4 mM DTT for another 10 min on ice. Pyruvate kinase activities were then assayed by measuring the amount of pyruvate production. Data are presented as the means ± SD (n=3 technical replicates). One-way ANOVA followed by post hoc test was used for the statistical analysis (\**p<*0.05; \*\*\**p<*0.001). (C) Upper: Glycerol gradient ultracentrifugation profiles of purified recombinant PKM2 and the effects of FBP and H_2_S on PKM2 oligomerization. Recombinant PKM2 (10 μg) was exposed to 100 μM FBP with or without 100 μM NaHS. After centrifugation, fractions were collected and analyzed by immunoblotting with PKM2 antibody. The distributions of PKM2 tetramer and dimer/monomer were as indicated. Bottom: Quantitative analysis of PKM2 protein level. The histogram represents normalized means ± SEM (n=3-5 biological replicates). The Student’s t-test was used for the statistical analysis (\**p<*0.05, group of FBP compared to the control group or group of FBP+NaHS). (D) Left: MDA-MB-231 cells were exposed to 1μM NaHS for 1hr. Subcellular localization of PKM2 was detected by immunocytochemistry. Nuclei were counterstained with DAPI. The representative images are shown. Scale bars: 10 μm. Right: The percentage of nuclear PKM2 is shown in a violin plot with individual points. Horizontal black dotted lines display the median and the percentage of cells with >20% nuclear localization of PKM2 were indicated (n= 65-70 individual cells, data were combined from three independent experiments). The Student’s t-test was used for the statistical analysis (\*\*\*\**p<*0.0001). (E) MDA-MB-231 cells were treated with 1μM NaHS for 24 hrs. The relative expression of genes regulated by PKM2 was measured by qRT-PCR. Data are presented as the means ± SD (n=3-5 biological replicates). The Student’s t-test was used for the statistical analysis (\**p<*0.05; \*\**p<*0.01; \*\*\**p<*0.001). (F) Cell proliferation assays were performed upon treatment with 0.25, 0.5 mM AOAA, or ddH_2_O control in MDA-MB-231 cells. Data are presented as the means ± SD (n=3 biological replicates). The Student’s t-test was used for the statistical analysis (\*\**p<*0.01, \*\*\**p<*0.001, 0.5 mM AOAA compared to the control group; ##*p<*0.01, ###*p<*0.001, 0.25 mM AOAA compared to the control group). (G) Western blot analysis of CBS, CTH, 3MST, and PKM2 expression in MCF-10A and MDA-MB-231 cells. GAPDH is the internal control. (H) Upper: MCF-10A and MDA-MB-231 cells were lysed and subjected to the biotin switch assay to precipitate sulfhydrated proteins. The biotin-labeled protein was analyzed by immunoblotting with anti-PKM2 antibody to detect sulfhydration of PKM2. Bottom: The SSH-labeled PKM2 was normalized with the level of total PKM2. Data are presented as the means ± SEM (n=4 biological replicates). The Student’s t-test was used for the statistical analysis (**p<0.01) (I) Upper: MDA-MB-231 cells were under normal (20% O_2_) or hypoxia incubator (1% O_2_) for 48h with or without 0.25 mM AOAA. Cells were then lysed and subjected to the biotin switch assay to precipitate sulfhydrated proteins. The biotin-labeled protein was analyzed by immunoblotting with anti-PKM2 antibody to detect sulfhydration of PKM2. Bottom: The SSH-labeled PKM2 was normalized with the level of total PKM2. Data are presented as the means ± SEM (n=4 biological replicates). One-way ANOVA followed by post hoc test was used for the statistical analysis (**p<0.01).

Studies have shown that PKM2 and two H_2_S-synthesizing enzymes, CBS and CTH, were upregulated in breast tumors ^22,27^. To further understand the clinical correlation between PKM2 and H_2_S-synthesizing enzymes in breast cancer, we analyzed expression profiles of PKM2, CBS, CTH, and 3-MST from the TCGA database. Upregulation of both PKM and CBS was observed in breast tumors (Fig. S2A), and CBS expression was associated with poor prognosis in breast cancer patients (Fig. S2B). Interestingly, correlation analysis reveals that expression of CBS is positively correlated with PKM, as well as most PKM2 response genes, including HIF1A, LDHA, GLUT1, GLUT12, and GLS1 (Fig. S2C), suggesting that elevated H_2_S levels in breast tumors, attributed by CBS upregulation, potentially enhance PKM2 dimerization and facilitate its translocation to the nucleus for transcriptional regulation. We then compared sulfhydration levels between mammary epithelial MCF-10A cells and breast cancer MDA-MB-231 cells. With higher CBS and CTH expression (Fig. 1G), significant higher sulfhydration levels were observed in MDA-MB-231 cells, as compared to MCF-10A (Fig. 1H). On the other hand, a previous study showed that excessive reactive oxygen species (ROS) induced by hypoxia can suppress the PK activity of PKM2 during cancer progression ^28^. Since cysteine sulfenylation (S-OH) by ROS is an important prior step for H_2_S to modify cysteine as protein sulfhydration ^29^, we assessed whether hypoxia can induce PKM2 sulfhydration in MDA-MB-231 cells. The level of PKM2 sulfhydration was increased under hypoxia, and this increment can be reduced by AOAA (Fig. 1I). Overall, our data suggest that in order to meet the increased biosynthetic demands in the hypoxic tumor microenvironment, cancer cells utilize H_2_S to stimulate PKM2 sulfhydration.

### Identification of the sulfhydration sites on PKM2 by mass spectrometry and mutation on cysteine 326 enhances the tetramerization of PKM2

There are ten cysteine residues located on each monomeric PKM2 (Fig. 2A). To further characterize the sulfhydration site on PKM2, we carried out liquid chromatography with tandem mass spectrometry (LC-MS/MS) on trypsin-digested PKM2 recombinant protein previously treated with NaHS (Fig. 2B) or endogenous PKM2 isolated from MDA-MB-231 cells (Fig. 2C). Results showed that PKM2 recombinant protein is sulfhydrated at cysteine 49 and 326 in the presence of NaHS (Fig. 2B). For endogenous PKM2 sulfhydration, to prevent excessive ROS-induced cysteine oxidation which may lead to artificial sulfhydration, we first added MMTS, which alkylates free thiol groups, into the culture medium before cell lysis. The clarified cell lysate was then subjected to immunoprecipitation to enrich PKM2 for sulfhydration analysis by LC-MS/MS. Only cysteine 326 on PKM2 was found to be sulfhydrated by this approach (Fig. 2C). We also checked whether other cysteine PTMs can be observed, including S-nitrosylation, glutathionylation, or oxidation. None of these PTMs was detected from our mass results (Table S1), although there may be other cysteine PTMs that cannot be detected by this proteomic method since most cysteine modifications are transient and reversible ^30^. Based on our proteomic analysis, cysteine 326 appears to be sulfhydrated endogenously in MDA-MB-231 cells (Fig. 2C). To further study the sulfhydration effects on cysteine 326, we substituted it with serine (PKM2^C326S^). Sulfhydration level was significantly reduced, but not eliminated, in the overexpressed HA-tagged PKM2^C326S^ in MDA-MB-231 cells (Fig. S3A), suggesting cysteine 326 acts as one of the major sulfhydration sites modulated by H_2_S. Consistent with the result we observed in cell lysates (Fig. S3A), there was less sulfhydrated recombinant PKM2 protein with C326S mutation, as compared to wildtype PKM2 (Fig. S3B). More importantly, by performing gel filtration analysis, we observed a striking transition from dimeric form to tetramer form in mutant PKM2^C326S^ recombinant proteins, while wildtype PKM2 recombinant proteins predominantly retained their dimeric form (Fig. 2D), suggesting PKM2 C326 may serve as a crucial allosteric site influencing PKM2 oligomerization. The elevated tetrameric configuration of PKM2 was also observed in cell lysates derived from MDA-MB-231 cells expressing the mutant PKM2^C326S^ (Fig. 2E), despite the complexity of cell lysates, which could encompass some PKM2 interacting proteins that may impact the distributions of PKM2.

**Figure 2.**
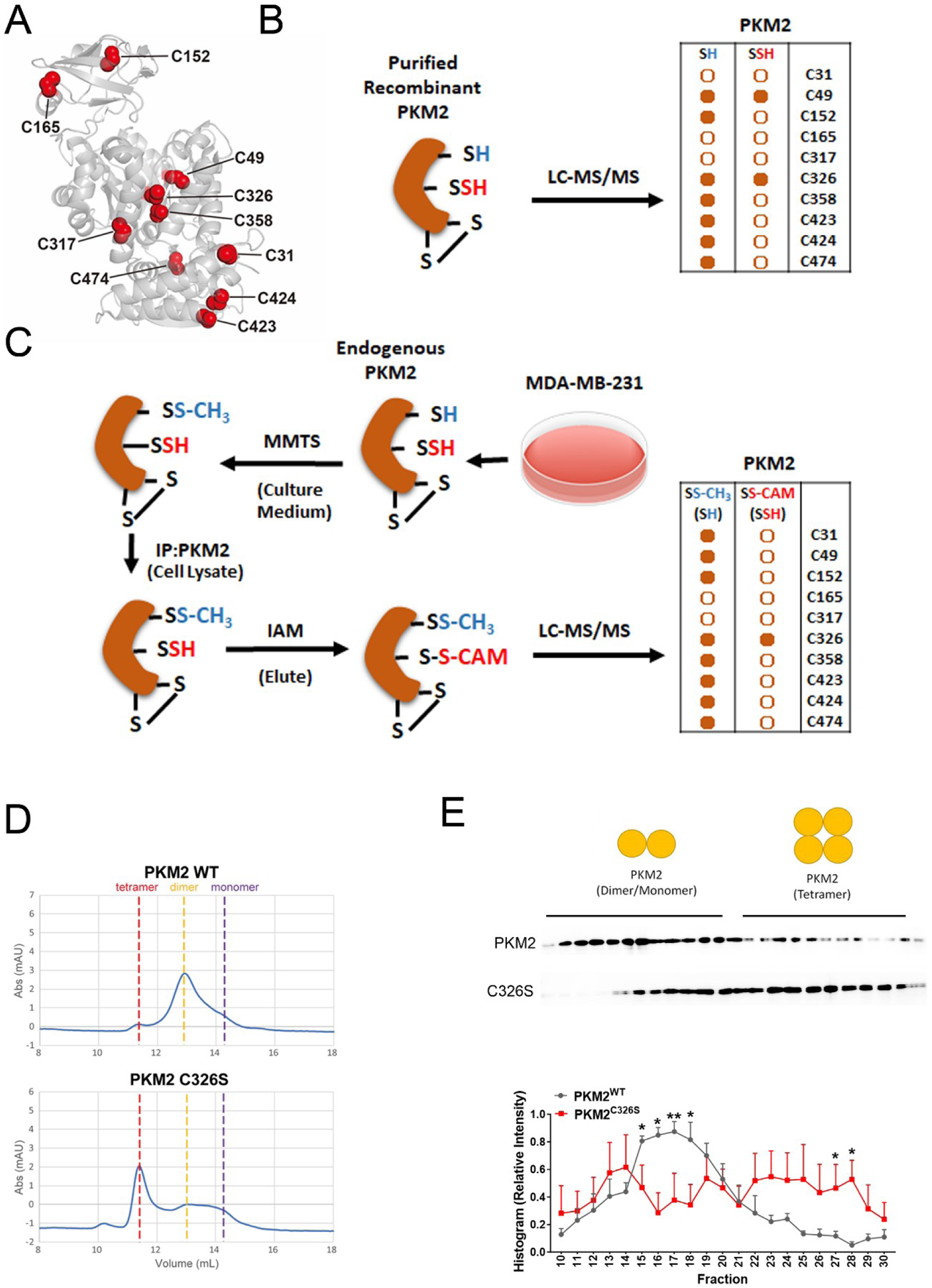
Replacement of PKM2 sulfhydration site to serine at C326 increases tetrameric oligomerization. (A) The PKM2 monomer shows the positions of ten cysteines in the folded structure. (B) Purified recombinant PKM2 was treated with 100 μM NaHS and then subjected to trypsin digestion. LC-MS/MS was performed. A side-by-side comparison of the thiol (-SH) and persulfide (-SSH) detected by MS on the same cysteine residues was shown. Dots with a solid orange color are depicted as positive detection by LC-MS/MS; the rest are depicted in hollow circles. (C) MDA-MB-231 cells were treated with 10 mM MMTS in the cell culture medium for 20 min. Cells were then lyzed and endogenous PKM2 was immunoprecipitated for LC-MS/MS analysis. A side-by-side comparison of the thiol (-SH) and persulfide (-SSH) detected by MS on the same cysteine residues is shown. Dots with a solid orange color are depicted as positive detection by LC-MS/MS; the rest are depicted in hollow circles. (D) Gel filtration analysis of PKM2 recombinant proteins (wildtype or C326S mutant) in the absence of FBP. The tetramer, dimer, and monomer are as indicated. (E) Upper: Glycerol gradient ultracentrifugation profiles of MDA-MB-231 cells transfected with V5-tagged PKM2 or PKM2^C326S^ mutant. After centrifugation, fractions were collected and analyzed by immunoblotting with anti-V5 antibody. Bottom: Quantitative analysis of PKM2 protein level. The histogram represents normalized means ± SEM (n=4 biological replicates). The Student’s t-test was used for the statistical analysis (\**p<*0.05; \*\**p<*0.01).

### Tetrameric PKM2^C326S^ adopts a unique conformation

PKM2 is allosterically regulated by the bound ligands. It is believed that tetrameric PKM2 adopts an inactive T-state conformation in the apo state or in the inhibitor-bound state and shifts to an active R-state conformation upon binding to FBP and allosteric activators ^31,32^. Each protomer consists of three functional domains (A, B, and C), with the active site between the A and B domains, and the FBP binding pocket in the C-terminal C domain. Two critical residues, W482 and W515, stabilize the T-state structure at the interface between two neighboring C domains (referred to as the C-C interface). FBP binding leads to global rotation of each protomer, disrupting the inter-molecular interaction of W482 and W515, while new inter-molecular contacts between K422 on one protomer and Y444 and P403 on the other are merged, giving rise to the fully active R-state. To delineate which tetrameric state the C326S mutant adopts, we determined the crystal structure of PKM2^C326S^ at 3.1 Å resolution in the apo form (Fig. 3A, Table S2). In one asymmetric unit, there are four protomers (A-D) belonging to two tetramers (AD/AD, BC/BC). These protomers have a folding similar to the wildtype (Fig. S4A-D). To provide the structural basis for the C326S mutation-mediated tetramerization, we compared the C326S to the T and R state conformers ^31,33^. Structural superimposition shows that PKM2^C326S^ is arranged in a different tetramer conformation (Fig. 3B-C).

**Figure 3.**
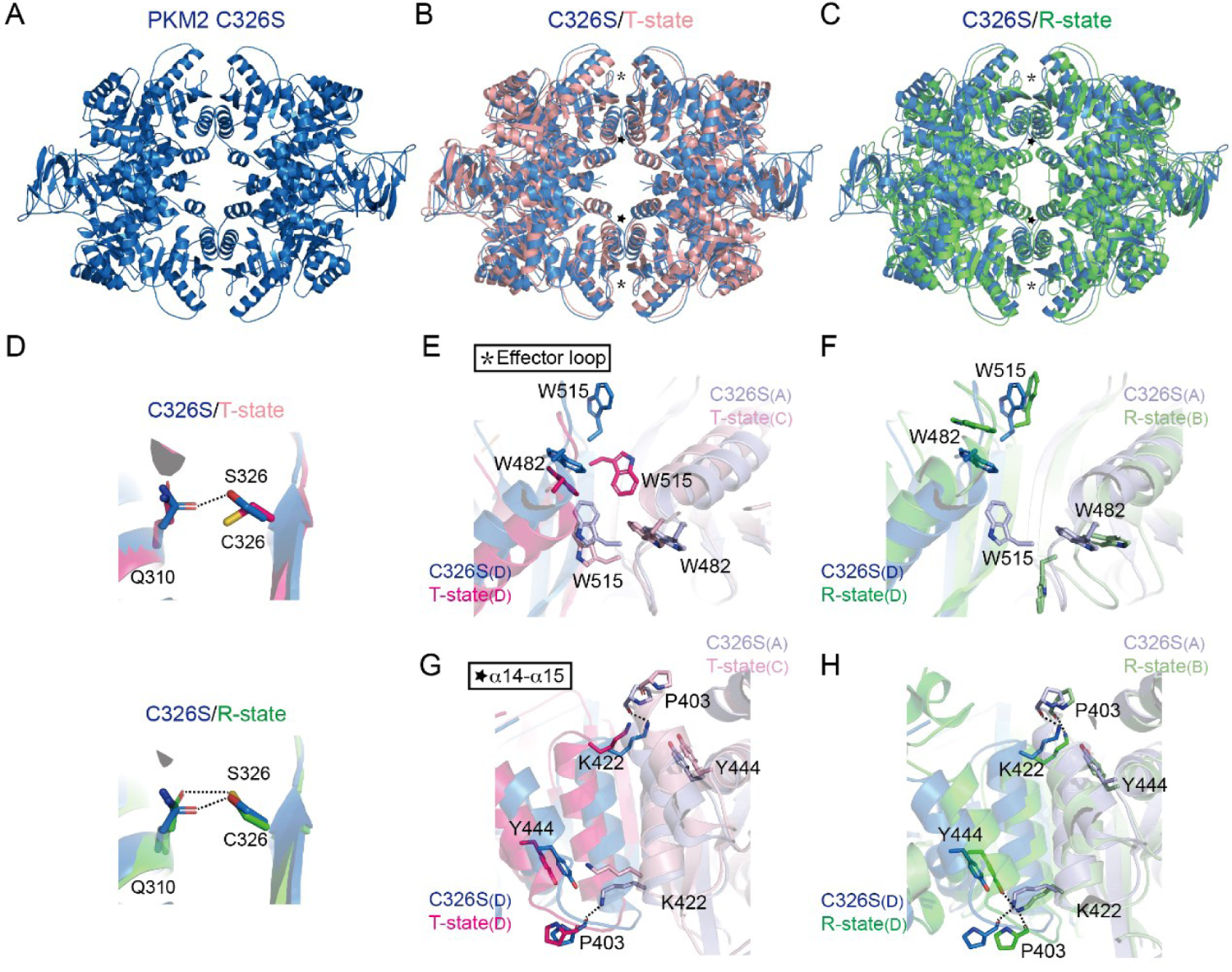
PKM2^C326S^ exhibits a unique conformation. (A) Crystal structure of the PKM2 C326S mutant. (B) Structural overlay of the C326S mutant on the T-state conformation (in complex with Phe, PDB: 4FXJ) ^31^. The effector loop and α14-α15 regions are indicated by asterisk and star respectively. (C) Structural overlay of PKM2^C326S^ on the R-state conformation (in complex with FBP and Ser, PDB: 4B2D) ^33^. PKM2^C326S^ is in blue, T-state in pink, and R-state in green. (D) Hydrogen bonding of residues 326 to Q310 in PKM2^C326S^ and the R-state conformation. (E, F) Zoom-in view of the effector loop region at the C-C interface. Side chains of W482 and W515 are shown in the stick presentation. Protomers are indicated in parentheses. (G, H) Zoom-in view of the α14-α15 region at the C-C interface. Hydrogen bonds are shown as dashed lines.

Located in the A domain, C326 makes contact with Q310 through hydrogen bonding in the R state, while this interaction is absent in the T state. In the C326S mutant, S326 also establishes hydrogen bonding with Q310 (Fig. 3D). This mutation does not change the overall conformation of the A domain. Instead, the C domains in PKM2^C326S^ display structural differences with an r.m.s.d. of 2.27 Å, while the C domains in the T or R state structure are similar with an r.m.s.d. of 0.96 Å and 0.64 Å, respectively (Fig. S4). The major differences come from the α14-α15 region and the effector loop, whose conformation is one of the key determinants for the T and R states. The AD/AD tetramer has better structural quality and is used for further structural analyses. Compared to the T-state conformation, the side chain of W515 from protomer D has swung away from the C-C interface, breaking the interaction between W515 (protomer D) and W482 (protomer A) (Fig. 3E-F). Additional inter-molecular contacts (cation-π interaction and hydrogen bonding) are established between Y444 and P403 on one protomer and K422 on the other in the α14-α15 region (Fig. 3G-H), which were observed in the R-state conformation. The abovementioned structural differences may result in the elevated tetramerization of the C326S mutant.

### Disruption of PKM2 sulfhydration at cysteine 326 promotes mitochondrial OXPHOS by increasing PK activity

To investigate the function of H_2_S-mediated PKM2 sulfhydration, we established breast cancer MDA-MB-231 cells and prostate cancer PC3 cells stably expressing PKM2^wt^ or PKM2^C326S^ mutant (Fig. 4A, S5A). The pyruvate kinase activities were significantly increased in MDA-MB-231 cells with PKM2^C326S^ expression (Fig. 4B). We next examined whether blocking PKM2 sulfhydration at C326 can rewire glucose metabolic fluxes. Remarkably, expression of PKM2^C326S^ significantly increased oxygen consumption rate (OCR) in both MDA-MB-231 cells (Fig. 4C, D) and PC3 cells (Fig. S5B, C), confirming our hypothesis that disruption of PKM2 sulfhydration redirects cancer cell glucose metabolism to mitochondrial oxidative phosphorylation (OXPHOS) system. The extracellular acidification rate (ECAR) was only slightly increased in MDA-MB-231 cells and PC3 cells expressing PKM2^C326S^ (Fig. 4E, F, S5D, E), suggesting the route to glycolysis was not much affected by the PKM2^C326S^ mutant. Restoring PK activities strongly facilitates OXPHOS, but not suppresses ECAR in both cell lines. To further characterize how PKM2 sulfhydration modulates glucose metabolism, we first checked expressions of PKM2-response genes, including CCND1, HIF-1α, LDHA, GLUT1, GLUT12, HK2, and GLS1. Most gene expressions were significantly reduced in MDA-MB-231 cells with PKM2^C326S^ expression, except CCND1 and LDHA (Fig. 4G). Metabolomic analysis of glycolytic intermediates was then performed using UPLC/Q-TOF-MS/MS. We found the levels of glucose, glucose-6-phosphate, fructose-1, 6-bisphosphate, ribose-5-phosphate, glyceraldehyde-3-phosphate, and phosphoenolpyruvate were significantly reduced in cells expressing PKM2^C326S^ mutant (Fig. 4H), revealing blockage of PKM2 sulfhydration at C326 leads to the increased PK activities to facilitate OXPHOS and results in the shortage of glycolytic intermediates and ribose-5-phosphate, which is essential for DNA synthesis. Meanwhile, the level of lactate was slightly increased in cells expressing PKM2^C326S^ compared to that in wildtype PKM2-expressing cells, while pyruvate was significantly upregulated in the C326S group (Fig. 4H). The changes of these metabolites were consistent with the observation from the seahorse, in which OCR was significantly increased in PKM2^C326S^ mutant while ECAR was only slightly increased (Fig. 4C-F). We next checked whether PKM2 sulfhydration at C326 is required for its nuclear translocation. As expected, the percentage of PKM2 level in the nucleus was much higher in the MDA-MB-231 cells expressing wildtype PKM2, as compared to the cells expressing PKM2^C326S^ (Fig. 4I). Overall, our data suggest that blocking PKM2 sulfhydration at C326 will facilitate mitochondrial OXPHOS by increasing PK activity and by suppressing its nuclear transcriptional activities (as illustrated in Fig. S5F).

**Figure 4.**
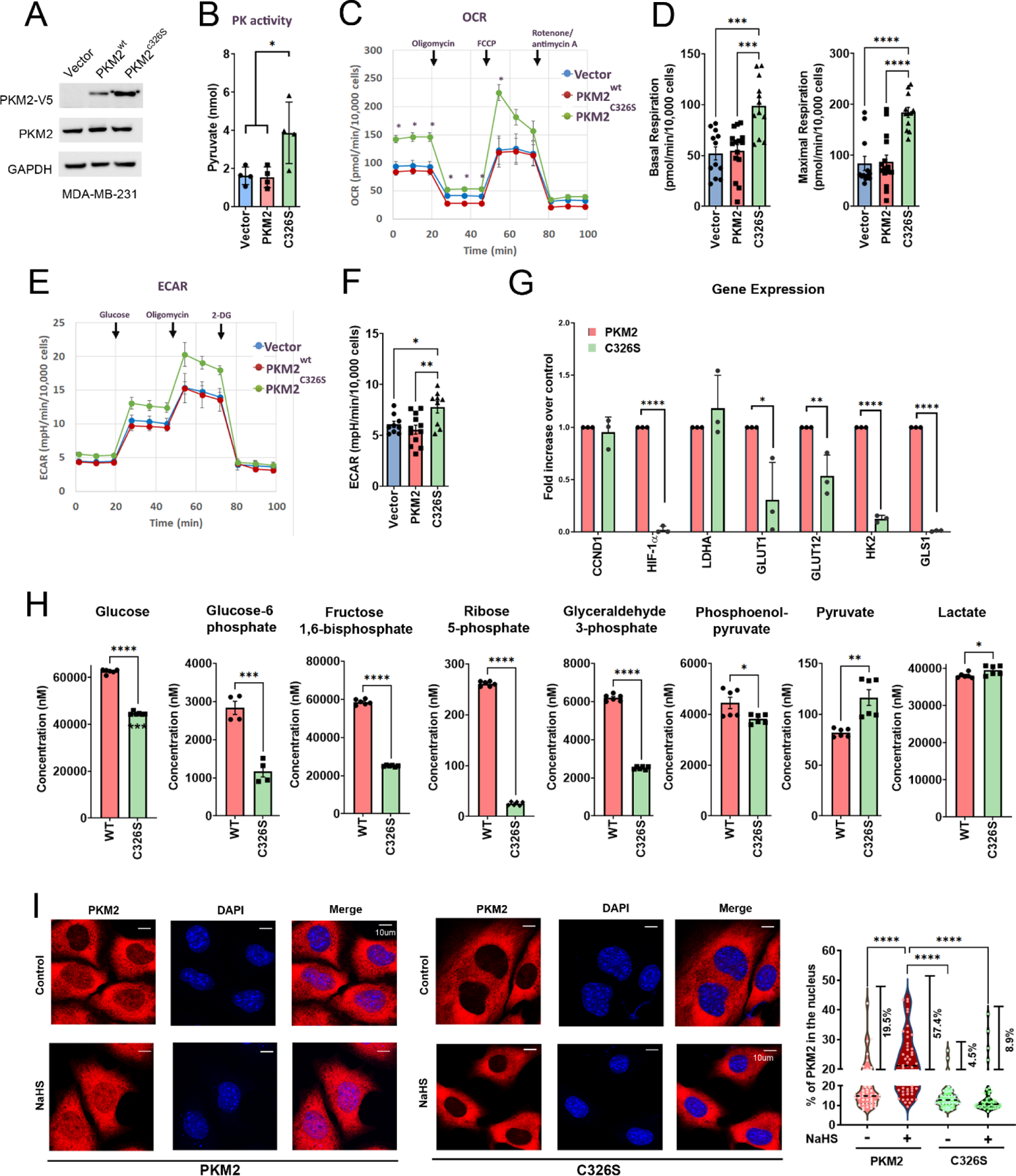
Blockage of PKM2 sulfhydration at C326 increases PK activity, resulting in enhanced mitochondrial oxidative phosphorylation and reduced nuclear translocation. (A) Western blot analysis of stable expression of V5-tagged PKM2 (wildtype or C326S mutant) in MDA-MB-231 cells. (B) Pyruvate kinase activities were assayed by measuring the amount of pyruvate production (nmol) in MDA-MB-231 cells expressing vector control, PKM2^wt^, or PKM2^C326S^ (n=3 biological replicates). One-way ANOVA followed by post hoc test was used for the statistical analysis (\**p<*0.05). (C) The oxygen consumption rate (OCR) curves in MDA-MB-231 expressing vector alone, PKM2^wt^, or PKM2^C326S^. Cells were treated with oligomycin, FCCP, and rotenone/antimycin A, respectively (n=3 biological replicates). The Student’s t-test was used for the statistical analysis (\**p<*0.05 C326S compared to the vector or PKM2 group). (D) The level of basal OCR and maximal OCR normalized to the cell numbers in MDA-MB-231 expressing vector alone, PKM2^wt^ or PKM2^C326S^ (n=9 or 12 experiments per group, data were combined from three independent experiments). One-way ANOVA followed by post hoc test was used for the statistical analysis (\*\*\**p<*0.001; \*\*\*\**p<*0.0001). (E) The ECAR curves in MDA-MB-231 expressing vector alone, PKM2^wt^ or PKM2^C326S^. Cells were treated with glucose, oligomycin, and 2-DG, respectively (n=3 biological replicates). (F) Glycolysis normalized to the cell numbers in MDA-MB-231 cells expressing vector alone, PKM2^wt^ or PKM2^C326S^ (n=9 or 12 experiments per group, data were combined from three independent experiments). One-way ANOVA followed by post hoc test was used for the statistical analysis (\**p<*0.05; \*\**p<*0.01). (G) The relative expression of PKM2-responded genes was measured by qRT-PCR from MDA-MB-231 cells with PKM2^wt^ or PKM2^C326S^. Data are presented as normalized means ± SD (n=3 biological replicates). Student t-test was used for the statistical analysis (\**p<*0.05; \*\**p<*0.01; \*\*\*\**p<*0.0001). (H) Metabolome analysis was conducted in MDA-MB-231 cells with PKM2^wt^ or PKM2^C326S^ expression. The level of glycolysis intermediates, including Glucose, Glucose-6-phosphate, Fructose-1,6-Bisphosphate, Ribose-5-phosphate, Glyceraldehyde-3-phosphate, Phosphoenolpyruvate, Pyruvate, and Lactate was determined by mass spectrometry. Data are presented as means ± SD (n=3-6 biological replicates). The Student’s t-test was used for the statistical analysis (\**p<*0.05; \*\**p<*0.01; \*\*\**p<*0.001, \*\*\*\**p<*0.0001). (I) Left: MDA-MB-231 cells expressing PKM2^wt^ or PKM2^C326S^ were exposed to 1μM NaHS for 1h. Subcellular localization of PKM2 was detected by immunocytochemistry. Nuclei were counterstained with DAPI. The representative images are shown. Scale bars: 10 μm. Right: The percentage of nuclear PKM2 is shown in a violin plot with individual points. Horizontal black dotted lines display the median and the percentage of cells with >20% nuclear localization of PKM2 were indicated (n= 41-47 individual cells, data were combined from three independent experiments). One-way ANOVA followed by post hoc test was used for the statistical analysis (\*\*\*\**p<*0.0001).

### PKM2 sulfhydration at cysteine 326 is required to facilitate cytokinesis during cell division

During cancer cell division, dimeric PKM2 with low PK activity facilitates biosynthesis from metabolic intermediates and exerts nucleus function to induce genes involved in the cell cycle ^34^, resulting in the promotion of G1/S cell cycle transition ^35^ and chromosome segregation during mitosis ^36^. To determine the effect of PKM2 sulfhydration on cell division, we first analyzed the level of cells in different phases of the cell cycle in MDA-MB-231 cells expressing wildtype PKM2 or PKM2^C326S^. The percentage of polyploid cells (>4N) was increased in cells expressing PKM2^C326S^ (Fig. 5A, B), suggesting blocking PKM2 sulfhydration at C326 may inhibit cytokinesis during mitosis. Interestingly, we also noticed that around 10% of the population of MDA-MB-231 cells with PKM2^C326S^ expression displayed giant shapes with multiple nuclei (Fig. 5C, D), which is one typical character of cell senescence ^37^. β-gal staining further confirmed that these giant cells were under senescence (Fig. S6A). Since dimeric PKM2 is known to play a pivotal role during metaphase and cytokinesis ^36,38^, we performed time-lapse microscopy at 10-min intervals continuously for 24 h to check whether disruption of PKM2 sulfhydration at C326 causes mitosis defects (Fig. 5E). Through observation following the mitotic progression in individual cells, we found that 22% of MDA-MB-231 cells with wildtype PKM2 showed cytokinesis failure (Fig. 5F), as judged by failure of separation of two daughter cells (Fig. 5E). Notably, cells expressing PKM2^C326S^ showed an approximately 1.5-fold increase in cytokinesis failure (Fig. 5F). Consistent with this observation, PKM2^C326S^ failed to interact with the spindle checkpoint protein BUB3 during metaphase (Fig. S6B), indicating that PKM2 sulfhydration is required to facilitate chromosome segregation, cytokinesis, and cell cycle progression. To confirm the increased cytokinesis failure in cells expressing PKM2^C326S^ is due to the inhibition of sulfhydration but not caused by other cysteine PTMs, we depleted H_2_S production by AOAA in MDA-MB-231 cells. Consistent with the observation from PKM2^C326S^ expressing cells, treatment with AOAA resulted in an increased percentage of polyploid cells (Fig. S6C-F). Overall, our findings indicate that blocking PKM2 sulfhydration at C326 or depletion of H_2_S by AOAA leads to impaired cytokinesis and subsequent inhibition of cell division.

**Figure 5.**
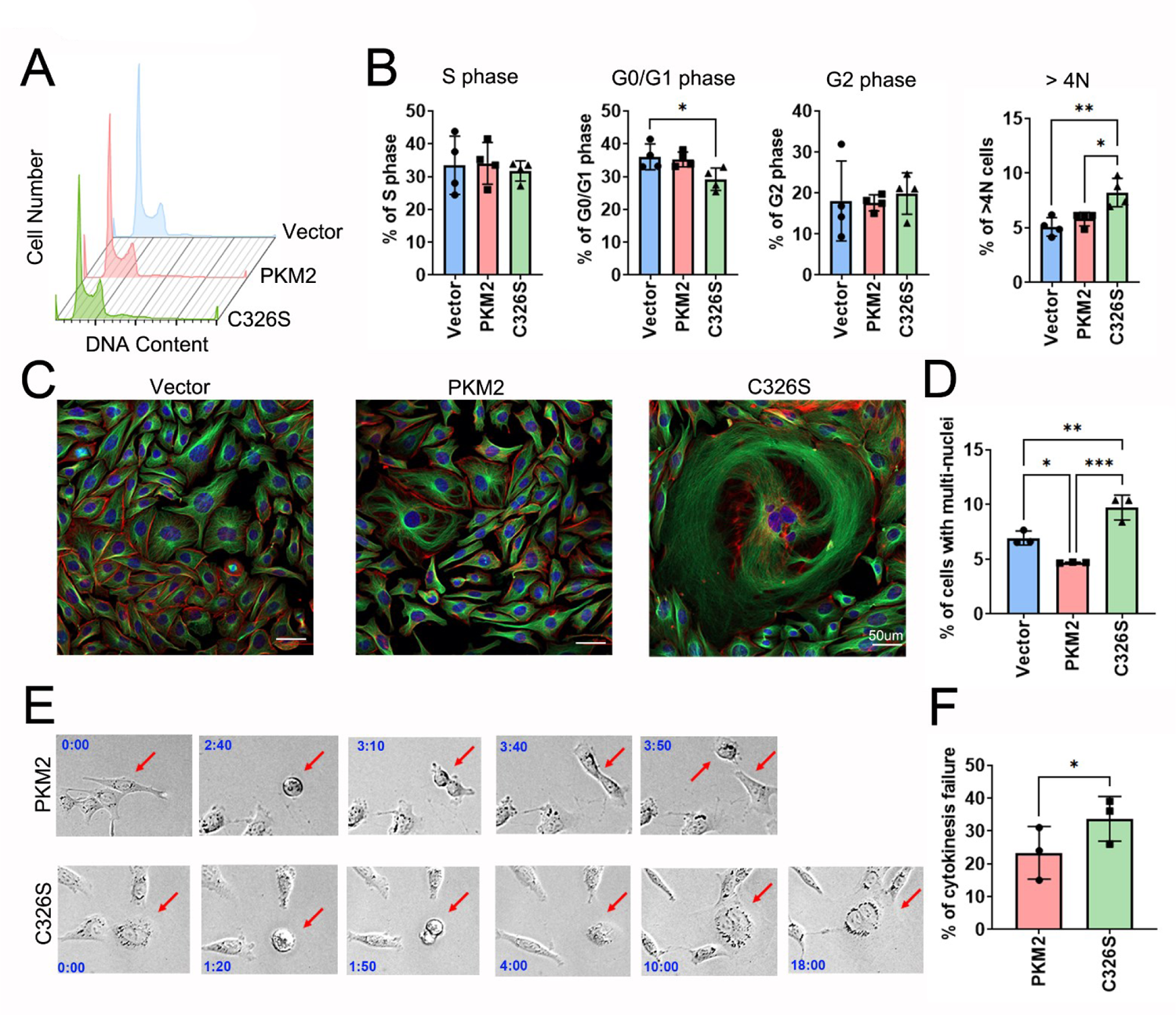
Blockage of PKM2 sulfhydration at C326 causes defects in cell cycle progression and cytokinesis failure. (A, B) The cell cycle of MDA-MB-231 cells expressing vector only, PKM2^wt^, or PKM2^C326S^ was examined by the PI staining method and measured by flow cytometry. (A) The representative image is shown. (B) The percentage of cells at different cell cycle phases was analyzed. Data are presented as means ± SD (n=4 biological replicates). One-way ANOVA followed by post hoc test was used for the statistical analysis (\**p<*0.05; \*\**p<*0.01). (C, D) Immunostaining of actin filaments (Red), α-tubulin (Green), and DNA counterstaining with DAPI (Blue) of MDA-MB-231 cells expressing vector only, PKM2^wt^, or PKM2^C326S^. (C) The representative images are shown. Scale Bar: 50 μm. (D) The percentage of poly-nuclei cells (n≧2) was analyzed. Data are presented as means ± SD (n=3 biological replicates). One-way ANOVA followed by post hoc test was used for the statistical analysis (\**P<*0.05; \*\**P<*0.01; \*\*\**P<*0.001). (E, F) Real-time live imaging of MDA-MB-231 cells expressing PKM2^wt^ or PKM2^C326S^ expression was performed to monitor cytokinesis failure. (E) Snapshots of a representative cell expressing PKM2^wt^ or PKM2^C326S^ undergoing division are shown. The red arrows point to cells undergoing mitotic division. Time stamps denote h:m of elapsed time. See also Movies S1 and S2 (F) Mitotic events with cytokinesis failure were recorded by time-lapse live cell microscopy. Results are expressed as means ± SD (n=3 biological replicates). The Student t-test was used for the statistical analysis (\**P<*0.05).

### Blockage of PKM2 sulfhydration reduces cell proliferation and inhibits tumor growth in the mouse xenograft model

Studies have shown that PKM2 tetramerization suppresses tumor growth by either overexpressing PKM1 or treatment with PKM2 activators ^11^. Consistently with this observation, the cell proliferation rate was dramatically reduced in MDA-MB-231 cells expressing PKM2^C326S^ mutant (Fig. 6A). More significantly, tumor growth was completely suppressed in the C326S group in the mouse xenograft model implanted with MDA-MB-231 cells expressing wildtype PKM2 or PKM2^C326S^ (Fig. 6B-F). Overall, our data demonstrated that disruption of PKM2 sulfhydration at C326 results in the significant suppression of tumor growth in the mouse xenograft model, and PKM2 sulfhydration may be an excellent therapeutic target for developing the anti-cancer drug.

**Figure 6.**
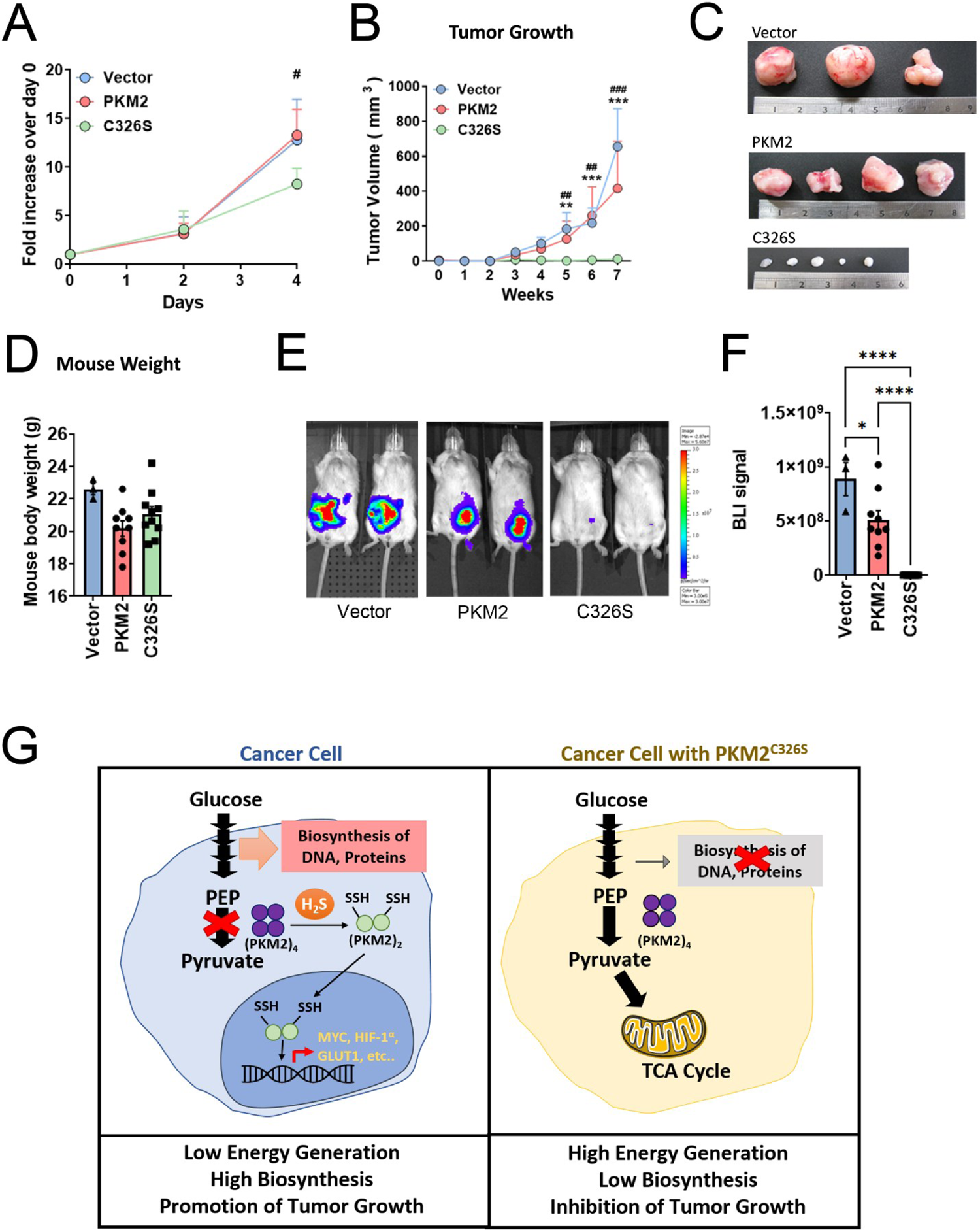
Blockage of PKM2 sulfhydration at C326 reduces cancer cell proliferation and tumor growth in the mouse xenograft model. (A) Comparison of cell proliferation rate between MDA-MB-231 cells expressing vector, PKM2^wt^, or PKM2^C326S^. Cell proliferation assay was performed and measured by MTS reagents. Data are the means ± SD (n=3 biological replicates). The Student’s t-test was used for the statistical analysis (#*p<*0.05, C326S compared to the PKM2 group). (B-F) MDA-MB-231 cells expressing vector, PKM2^wt^ or PKM2^C326S^ were injected into the mouse mammary fat pad. (B) The tumor volumes were monitored every week. Data are means ± SD (n=3-10; data were combined from two independent experiments). The Student’s t-test was used for the statistical analysis (\*\**p<*0.01, \*\*\**p<*0.001, C326S compared to the vector group; ##*p<*0.01, ###*p<*0.001, C326S compared to the PKM2 group). (C) Representative pictures of tumors of each group. (D) The mean of mouse weight ± SD (n=3-10; data were combined from two independent experiments) (E) Representative bioluminescence images (BLI) at week 9 after implantation of respective cancer cells are shown. (F) The kinetics of individual tumors was monitored by IVIS. Data are means ± SD (n=3-10; data were combined from two independent experiments). One-way ANOVA followed by post hoc test was used for the statistical analysis (\**p<*0.05; \*\*\*\**p<*0.0001). (G) A schematic illustration revealing H_2_S modulates glucose metabolism switch through destabilizing PKM2 oligomerization. Left: H_2_S promotes dissociation of PKM2 tetramer to dimer through protein sulfhydration. Inhibition of PKM2 activity results in the accumulation of metabolic intermediates required for the biosynthesis. Meanwhile, the PKM2 dimers translocate into the nucleus to facilitate multiple gene expressions to promote cancer progression. Right: Blockage of PKM2 sulfhydration at cysteine 326 by mutation stabilizes PKM2 tetramer to maintain high PK activity, resulting in high energy generation, low biosynthesis, and inhibition of tumor growth.

## Discussion

Metabolic alteration is one of the major hallmarks of cancer. In order to grow rapidly, tumor cells display notable increases in glucose uptake and higher demand on metabolic intermediates for biosynthesis. In this study, we show that H_2_S mediates PKM2 activities through protein sulfhydration, resulting in the dissociation of the high-activity tetramer form to low low-activity dimer form. This PKM2 modification rewires glucose metabolism to meet demands on high biosynthesis and low energy production for cancer cells to proliferate rapidly. In contrast, blocking PKM2 sulfhydration at C326 by mutation or treatment of H_2_S inhibitor, AOAA, prevents tetramer dissociation and nuclear translocation of PKM2, facilitating oxidative phosphorylation for high energy production and low biosynthesis, leading to the significant inhibition of tumor growth. Moreover, PKM2^C326S^ forms heterocomplexes with wildtype PKM2, resulting in significantly enhanced PK activity, suggesting C326 is a potential target for developing small molecule drugs to target cancer metabolism (Fig. 6G).

Cysteine is the major target for redox PTMs in response to environmental stresses ^30^. In this study, we investigated the significance of cysteine sulfhydrations on PKM2 oligomerization, particularly focusing on cysteine 326. Using a proteomic approach, we demonstrated that cysteine 326 was sulfhydrated endogenously (Fig. 2C). In the meantime, none of the other cysteine PTMs was detected at cysteine 326 from our LC-MS/MS analysis (Table S1). Despite it is challenging to rule out the possibility of other PTMs occurring at this site since most cysteine modifications are transient and reversible ^30^. In fact, these modifications can sometimes occur sequentially; for instance, H_2_S can react with oxidized, S-nitrosated cysteines, or cysteine disulfides, to form sulfhydrated cysteines ^22,29,39^. To assess the importance of sulfhydration specifically and distinguish it from other modifications, we substituted cysteine with serine at position 326. However, this substitution not only blocked sulfhydration but also prevented other potential modifications. To unravel whether the significance we observed in PKM2^C326S^ is majorly due to sulfhydration, we examined PKM2 functionality in cells with H_2_S depletion achieved by AOAA treatment. Consistent with the results obtained from PKM2^C326S^ (Fig. 4-6), treatment with AOAA led to the inhibition of PKM2 nuclear localization (Fig. S1D), cytokinesis inhibition (Fig. S6C-F), as well as cell proliferation (Fig. 1F). Overall, our data obtained from AOAA treatment support the hypothesis that the effects observed upon blocking modifications at cysteine 326 of PKM2 are predominantly mediated through sulfhydration.

Structurally, replacing the Cys with Ser at residue 326 of PKM2 remotely changes the dynamics of the C domain and promotes tetramerization (Fig. 3, S4). Considering the importance of PKM2 in cancer metabolism, PKM2 has emerged as a therapeutic target in treating cancer. Small-molecule inhibitors that selectively activate PKM2 have been developed, such as TEPP-46 ^40,41^. These synthetic PKM2 activators promote the association of PKM2 subunits into constitutive tetramers, resulting in increased enzymatic activities and reduced tumor size in mouse xenografts ^11^. Here our study reveals a novel regulatory mechanism on PKM2 activity, and PKM2^C326S^ mutation that blocks sulfhydration results in a higher amount of tetrameric state in the absence of FBP or other activators (Fig. 2D, E). Interestingly, the tetramer organization of PKM2^C326S^ is different from the currently known T or R states. Compared to all known PKM2 activators which activate PK activity constitutively, activators targeting PKM2^C326S^ may activate PK activity in a much more moderate way, which may lower the undesired side effects on normal cells for their growth and function.

Cell cycle checkpoints are the surveillance mechanisms that ensure the proper progression of cell division ^42^. Among them, the spindle assembly checkpoint (SAC) at the metaphase-anaphase transition is essential for the checkpoint delaying anaphase until the last chromosome is aligned on the metaphase plate ^43^. It has been reported that PKM2 interacts with and phosphorylates Bub3 in metaphase, which is important for the recruitment of the Bub3-Bub1 complex to the Blinkin and the microtubule spindle attachment to kinetochores ^36^. Here our data indicate PKM2^C326S^ mutant failed to bind Bub3 in metaphase (Fig. S6B), which may lead to the misalignment of chromosomes, and subsequently, cause abnormal chromosome segregation and lagging chromatids in anaphase ^36^. This condition is called aneuploidy, which results in high-level micronuclei and may further cause cytokinesis failure ^44^, as we observed in cells expressing PKM2^C326S^ mutant (Fig. 5), as well as in cells with H_2_S depletion (Fig. S6C-F).

Most cancer cells reprogram their metabolism in favor of aerobic glycolysis to meet their bioenergetics and biosynthetic demands during tumorigenesis. In breast cancer, one of the H_2_S-producing enzymes, CBS, was upregulated in breast cancer cells and clinical tumor tissues, as compared to normal cells and normal tissues, respectively (Fig. 1G, S2A). The elevated levels of CBS were positively associated with increased sulfhydrated PKM2 levels in a breast cancer MDA-MB-231 cell line (Fig. 1H), along with upregulated expressions of PKM2-response genes in clinical samples (Fig. S2C), suggesting cancer cells rewire glucose metabolism through modifying PKM2 activity by H_2_S-mediated sulfhydration. On the other hand, the PKM2 sulfhydration was elevated in breast cancer cells under hypoxia (Fig. 1I), suggesting cancer cells may alter their metabolic pathways through PKM2 sulfhydration in response to the hypoxic tumor microenvironment. By blocking H_2_S mediated sulfhydration at C326, we provided a promising therapeutic opportunity in which we activate PKM2, resulting in glucose metabolic switch (Fig. 4) and complete tumor growth inhibition in the mouse xenograft model (Fig. 6). Previous study indicates that the expression of glucose transporters, such as GLUT1, and glucose metabolic enzymes are high in breast cancer, especially in triple-negative breast cancer (TNBC) ^45^, suggesting glucose metabolism may act as the main source for energy and building materials in breast cancer. Moreover, glucose metabolism is associated with tumor growth, metastasis, and drug resistance in breast cancer ^46–48^. Therefore, in our study, breast cancer responds significantly when we target glucose metabolism through the blockage of H_2_S-mediated PKM2 sulfhydration. Developing inhibitors that block PKM2 sulfhydration can be considered a metabolic therapy for breast cancer patients.

In addition to the modulation of cancer glucose metabolism, endogenously produced H_2_S is also essential and sufficient for dietary restriction extended life span ^20^. The following study further indicates that H_2_S-induced angiogenesis under amino acid restriction is through inhibition of mitochondrial OXPHO, increased glycolytic ATP production, and increased glucose uptake in endothelial cells ^49^. Here our data further support their discovery and provide an excellent explanation of the molecular basis that H_2_S may facilitate the OXPHO/glycolysis switch by destabilizing PKM2 tetramerization. We also should take PKM2 sulfhydration into consideration on other known H_2_S mediated functions, such as vasodilation, immunomodulation, or pro-survival signals to decipher their underlying mechanisms under normal physiological conditions or dysregulated disease conditions.

## Methods

### Antibodies

PKM2 (D78A4) XP® Rabbit mAb (#4053), HA-Tag (C29F4) Rabbit mAb (#3724), and V5-Tag (D3H8Q) Rabbit mAb (#13202) were from Cell Signaling Technology (CST). CTH Antibody (F-1) (#sc-374249), GAPDH Antibody (6C5) (#sc-32233), Tubulin Antibody (B-5-1-2) (#sc-23948), mouse anti-rabbit IgG-HRP (#sc-2357), and m-IgGκ BP-HRP (anti-mouse) (#sc-516102) were from Santa Cruz Biotechnology. Goat anti-Rabbit IgG (H+L) Cross-Adsorbed Secondary Antibody, Alexa Fluor™ 594 (#A-11012) and Goat anti-Mouse IgG (H+L) Cross-Adsorbed Secondary Antibody, Alexa Fluor™ 488 (#A-11001) were from Invitrogen. Recombinant Anti-CBS antibody [EPR8579] (#ab140600) and Recombinant Anti-Bub3 antibody [EPR5319(2)] (#ab133699) were from abcam. MPST antibody [C2C3], C-term (#GTX108274) was from GeneTex. Rabbit IgG, Purified (#PP64B) was from EMD Millipore.

### Chemicals

S-Methyl methanethiosulfonate (MMTS) (#64306), Iodoacetamide (IAM) (#I6125), and O-(Carboxymethyl)hydroxylamine hemihydrochloride (AOAA) (#C13408) were from Sigma-Aldrich.

### Cell culture and transfection

MDA-MB-231 and PC3 cell lines were cultured in Dulbecco’s Modified Eagle’s Medium (DMEM, Gibco) supplemented with 10% fetal bovine serum (FBS, HyClone) and 1% Penicillin/Streptomycin (P/S, Gibco). MDA-MB-231 and PC3 cells stably expressing GFP-Luciferase, together with pLX304, pLX304-PKM2, or pLX304-PKM2-C326S, were generated by lentivirus infection and established as mass culture, selected and maintained in 1 μg/mL puromycin (Cyrusbioscience, #101-58-58-2) and 1 μg/mL Blasticidine S hydrochloride (BSD, Cyrusbioscience, #101-3513-03-9), respectively. MCF-10A cell line was cultured in DMEM/F-12 (Gibco) supplemented with 5% horse serum (HyClone), 20 ng/mL epidermal growth factor (EGF, PEPROTECH, #AF-100-15), 0.5 μg/mL hydrocortisone (Sigma-Aldrich, #H0888), 10 μg/mL insulin (Sigma-Aldrich, #I9278), and 1% Penicillin/Streptomycin (P/S, Gibco). All cells were cultured at 37 °C and 5% CO_2_ with humidity. Plasmids were transfected using Opti-MEM (Gibco, #31985070) and TransIT-X2 (Mirus, #MIR 6000).

### Immunocytochemistry (ICC)

Cells (0.8-3 × 10^4^ cells) were seeded on poly-L-lysine (0.01%, Sigma-Aldrich, #P8920) and fibronectin (10 μg/mL, Sigma-Aldrich, #FC010) coated glass coverslips overnight. For treatment with NaHS, the cell culture medium was changed to a serum-free medium and incubated in a standard incubator. After 24-hour incubation, cells were treated with 1μM NaHS for 1 hour. For treatment with AOAA, cells were treated with 0.25 mM AOAA or H_2_O control for 24 to 72 hours. Then, cells were fixed in 4% formaldehyde for 30 min, washed three times with PBS, and permeabilized with 0.1% Triton X-100 in PBS for 10 min. Cells were incubated in blocking buffer (1% BSA in PBS with 5% FBS, filtered before use) for 1 hour at room temperature before incubation with Rhodamine Phalloidin (Invitrogen, #R415), primary [PKM2 (D78A4) XP® Rabbit mAb, or Anti-α Tubulin Antibody (B-5-1-2)] and secondary [Goat anti-Rabbit IgG (H+L) Cross-Adsorbed Secondary Antibody, Alexa Fluor 594 or Goat anti-Mouse IgG (H+L) Cross-Adsorbed Secondary Antibody, Alexa Fluor™ 488] antibodies. Incubations with both primary and secondary antibodies (diluted in PBST) were done in the presence of 1% FBS. Mounting medium (EverBrite, #23008-T) with 1 μg/mL DAPI (Sigma-Aldrich, #D8417) was used to adhere a coverslip to a slide glass. Fluorescence was visualized using a confocal laser scanning microscope (ZEISS LSM 800) or fluorescence microscope (Nikon ECLIPSE Ni microscope). Images were analyzed by ZEN 2012 (Carl Zeiss) and ImageJ ^50^ software.

### Time-lapse live cell microscopy

Cells were seeded in 35 mm diameter dishes (Alpha Plus; Scientific Corp., Taoyuan, Taiwan) overnight and imaged with a Leica DMI6000 microscope (Leica Microsystems, Wetzlar, Germany) equipped with an HCX PL FL × 20/0.4 numerical aperture objective and an Andor Luca R EMCCD camera (Andor Technology, Belfast, United Kingdom). Differential interference contrast images from multiple positions were acquired at a 10-minute interval for 24 hours. The percentage of cells that undergo cytokinesis failure was counted manually. At least 40 cells undergoing mitosis were counted for each experiment and repeated independently three times. All images were analyzed with MetaMorph software (Molecular Devices, San Jose, CA, USA).

### Quantitative real-time PCR

Cells were treated with serum-free medium containing 10 µM NaHS or growth medium containing 0.25 mM AOAA for 24 hours before RNA extraction. RNA was extracted from cells using TRIzol (Invitrogen, #15596018) and Chloroform Replacement (Cyrusbioscience, #CRSR) following protocols supplied by the manufacturer. First-strand cDNA was generated by ReverTra Ace (PURIGO, #PU-TRT-100) using oligo-dT. Real-time PCR was performed on a qTOWER³ real-time PCR detection system (Analytik Jena) and analyzed with qPCRsoft version 3.4 software (Analytik Jena). The KAPA SYBR® FAST qPCR Master Mix (2X) Kit (Roche, KK4600) was used. The mRNA levels were normalized to that of actin or GAPDH. All primer sequences used in this study are listed in Table S3.

### Modified biotin switch (S-sulfhydration) assay

The assay was performed following a published paper, with modifications ^51^. Briefly, after different treatment conditions, cells were homogenized in a HEN buffer [250 mM HEPES-NaOH (pH 7.7, Sigma-Aldrich, #H4034)), 1 mM EDTA, and 0.1 mM neocuproine], supplemented with 100 μM deferoxamine and 1X protease inhibitor (Roche), and centrifuged at 13,000g for 30 min at 4 °C. Cell lysates (0.4-1 mg) or recombinant proteins (10-50 μg) were treated with 100 µM NaHS at 37 °C for 30 min and then 1 mM DTT for another 10 or 30 min at 37 °C if needed. The lysates were then added to a blocking buffer (HEN buffer adjusted to 2.5% SDS and 20 mM MMTS) at 50 °C for 20 min with frequent vortexing. The proteins were precipitated by 100% acetone at −80 °C for at least 20 min. After acetone removal, the proteins were resuspended in a HENS buffer (HEN buffer adjusted to 1% SDS), and then 4 mM biotin-HPDP (EZ-link, ThermoFisher) was added. After incubation for 3 hours at RT, biotinylated proteins were precipitated by streptavidin-agarose beads at 4 °C overnight and then washed with the wash buffer [25 mM HEPES (pH 7.5), 1 mM EDTA, 600 mM NaCl, and 0.5% Triton X-100]. The biotinylated proteins were analyzed by SDS-polyacrylamide gel electrophoresis and subjected to Western blot analysis.

### Western blotting

Cells were lysed in a 1X RIPA buffer [50 mM Tris buffer, pH 7.4, 150 mM NaCl, 1% Triton X-100, 1 mM EDTA, 0.1% SDS, and 1X protease inhibitor (Roche)]. The cell lysates were resolved in 10 or 12.5% SDS-polyacrylamide gel, transferred onto the PVDF membrane, and probed with antibodies. ECL reagent (Perkin Elmer, #NEL105001EA or Thermo Scientific, #34095) was used to capture luminescence by ImageQuant™ LAS 4000 (GE).

### Real-time measurements of OCR and ECAR

Seahorse XFe24 Analyzer (Agilent) was used for cell mito stress test and glycolysis stress test, i.e., measured oxygen consumption rate (OCR) and extracellular acidification rate (ECAR). Seahorse XFe Wave software (Agilent) was used for data analysis and report generation. Cells were seeded into the wells of the microplate at different densities, which would be a monolayer on the next day, in 100 μL of culture medium. After 1-5 hours, cells were attached and 150 μL of culture medium was added. On the day of assay, culture media were removed gently, assay media (DMEM, without buffer reagent, 2% FBS) were added, and cells were incubated at 37 °C in the absence of CO_2_.

Cell Mito Stress Test (Agilent, #103015-100):

The test was performed according to the methods described in the manufacturer manual. Basal OCR was measured three times before adding 1 μM oligomycin (inhibitor of ATP synthase). The second injection was carbonyl cyanide-4 (trifluoromethoxy) phenylhydrazone (FCCP), which is an uncoupling agent that collapses the proton gradient and disrupts the mitochondrial membrane potential. The third injection is a mixture of rotenone (inhibitor of complex I) and antimycin A (inhibitor of complex III).

### Glycolysis Stress Test (Agilent, #103020-100)

The test was performed according to the methods described in the manufacturer manual. ECAR of cells was measured three times before and after sequential injections of three different compounds: 10 mM glucose, 1 μM oligomycin (inhibitor of ATP synthase), and 100 mM 2-Deoxy-D-glucose (2-DG, a glucose analog that inhibits glycolysis).

### Purification of recombinant proteins

His-tagged WT and mutant PKM2 were expressed in *Escherichia coli* incubated at 37 °C and inducted with 0.1 mM isopropylthiogalactopyranoside (IPTG) at 25°C for 5 hours. Recombinant proteins were then purified by Ni-NTA-agarose (Qiagen) affinity chromatography. The elution buffer contained 20 mM Tris (pH 8.0, APOLO Biochemical, #APL-0082), 150 mM potassium chloride (KCl, Sigma-Aldrich, #P5405), 1 mM β-mercaptoethanol (Cyrusbioscience, #101-60-24-2), 1 mM PMSF, and 100-400 mM imidazole (Cyrusbioscience, #101-288-32-4). The eluate proteins were further purified by gel filtration chromatography (Superdex 200 Increase 10/300 GL, Cytiva) in PBS buffer (pH 8.0) and 10% glycerol (J.T. Baker, #2136-08). Protein concentration was determined by the absorbance at 280 nm by a Colibri microvolume Spectrophotometer (Titertek-Berthold). All proteins were flash-frozen by liquid nitrogen and stored as aliquots at −80 °C.

### Pyruvate kinase assay

The pyruvate kinase reaction rate was determined by LDH-coupled enzyme assays. In a 200 μL PBS (pH 8.0) buffer containing 5 mM MgCl_2_ (SHOWA chemical, #1301-7260), 1 mM ADP (Sigma-Aldrich, #A2754), 0.4 mM NADH (Sigma-Aldrich, #N8129), 2 U LDH (Sigma-Aldrich, #L1254), and cell lysates or PKM proteins were incubated at room temperature for 5 min. The addition of 0.5 mM PEP (Sigma-Aldrich, #P0564) initiated the PKM-LDH coupled reaction.

The absorbance at 340 nm was measured every 10 or 12 seconds for 15 min by a Synergy HTX plate reader (BioTek). An NADH standard curve was used to determine the reaction rate. All reported rates were normalized by protein concentration. NaHS was incubated with protein on ice for 30 min and then assayed immediately. FBP was from Sigma-Aldrich (#F6803).

### Mass spectrometry analysis to detect PTM in recombinant proteins

His-tagged recombinant PKM2 proteins were treated with 100 μM NaHS for 30 min at 37 °C and then performed in solution trypsin digestion. Peptide digests are further purified by C18 Zip-tip and evaporated in a speed vacuum before injection. Samples were then dissolved in 8 μL of 2% acetonitrile with 0.1% trifluoroacetic acid. Samples were injected into the Waters NanoAcquity UPLC-Synapt G2 HDMS at National Health Research Institutes mass spectrometry facilities. The sulfhydrated modification on PKM2 was further analyzed by MASCOT.

### Mass spectrometry analysis to detect endogenous PTM in cell lysates

#### On-Bead Digestion

MDA-MB-231 cells were pre-incubated with 100 nM NaHS for 30 min followed by 10 mM MMTS for 20 min before lysed in non-denaturing lysis buffer [20 mM Tris-HCl pH 8, 137 mM NaCl, 1% NP-40, 2 mM EDTA, and 1X protease inhibitor (Roche)]. The cell lysates were then centrifuged, and the supernatants were collected and diluted to the same concentration. The cell lysates were then mixed with anti-PKM2 antibody (CST) and Protein A Mag Sepharose Xtra (Cytiva). The mixture was gently rocked overnight at 4 °C on the rocker. The next day, the beads were washed 3 times with the non-denaturing lysis buffer and then 3 times with PBS. Then, the beads were suspended in 50 mM HEPES (pH 8) with 25 mM IAM at RT for 30 min. Afterward, samples were sequentially digested by Lys-C protease (FUJIFILM Wako Pure Chemical Corporation, #125-05061) at 37 °C for 3 hours, followed by trypsin (Promega, #V511A) at 37°C overnight. After digestion, the peptide mixtures were extracted and desalted using Pierce C18 Spin Tips (Thermo Scientific, #84850) according to the manufacturer’s protocol. Elutes were dried under vacuum and re-suspended in 0.1% formic acid (J.T. Backer, #9834-03).

#### Shotgun Proteomic Identifications

NanoLC-nanoESI-MS/MS analysis was performed on a nanoAcquity system (Waters, Milford, MA) connected to the Orbitrap Elite hybrid mass spectrometer (Thermo Electron, Bremen, Germany) equipped with a PicoView nanospray interface (New Objective, Woburn, MA). Peptide mixtures were loaded onto a 75 μm ID, 25 cm length C18 BEH column (Waters, Milford, MA) packed with 1.7 μm particles with a pore of 130 Å and were separated using a segmented gradient in 30 min from 5% to 25% (0 min to 27.5 min), 25% to 35% (28.5 min to 30 min) solvent B [acetonitrile (J.T. Backer, #9829-03) with 0.1% formic acid] at a flow rate of 300 nL/min and a column temperature of 35 °C. Solvent A was 0.1% formic acid in water. The mass spectrometer was operated in the data-dependent mode. Briefly, survey full scan MS spectra were acquired in the orbitrap (m/z 350-1600) with the resolution set to 60K at m/z 400 and automatic gain control (AGC) target at 106. The 15 most intense ions were sequentially isolated for HCD MS/MS fragmentation and detection in the orbitrap with previously selected ions dynamically excluded for 60s. For MS/MS, we used a resolution of 15000, an isolation window of 2 m/z, and a target value of 50000 ions, with maximum accumulation times of 200 ms. Fragmentation was performed with a normalized collision energy of 30% and an activation time of 0.1 ms. Ions with single and unrecognized charge states were also excluded. For targeted LC-MS/MS experiments, inclusion list mass tolerances were set to ±10 ppm, and monoisotopic precursor ion selection was enabled.

#### Database searching

Raw data files were processed using Proteome Discoverer v3.0.1.27 (Thermo Fisher Scientific) and the tandem MS data were then searched using SEQUEST algorithms against the protein sequence of PKM2, Protein A, Ig gamma chain C region, and a human UniProt (Swiss-Prot only) database (released February 2023) with common contaminant proteins. The search parameters included trypsin as the protease with a maximum of 2 missed cleavages allowed; +15.9949 Da (oxidation of methionine and cysteine), +0.984 Da (deamidation of asparagine and glutamine), +31.990 Da (dioxidation of cysteine), +47.985 Da (trioxidation of cysteine), +57.021 Da (carbamidomethylation of cysteine), and +88.994 Da (C2H3NOS, Cys-S-S-CAM, for IAM-tagged persulfides) were set as a dynamic modification. Precursor mass tolerance was set to 10 ppm and fragment mass tolerance was set to 0.02 Da. The false discovery rates of proteins and peptides were set to 0.01. These data have been deposited in the PRoteomics IDEntifications (PRIDE) database (Project accession: PXD044925).

### Mass spectrometry analysis for quantitation of metabolites

Cell pellets were re-suspended in 1 mL 100% methanol (precooled to −80 °C) and quenched in liquid nitrogen. The frozen-quenched cells were thawed, vortexed for 30-60 seconds, and pelleted by centrifugation. The supernatant was transferred to a fresh tube, the cell pellets were re-suspended in 1 mL 100% methanol (precooled to −80 °C), and the following procedures were repeated. The supernatant was evaporated in a centrifugal evaporator to generate a dry metabolite pellet. Xevo TQ-S triple quadrupole (tandem quadrupole) mass spectrometer (Waters) was used for the quantitation of metabolites involved in the pathways of glycolysis and the TCA cycle.

### Cell proliferation assay

The cell proliferation assay was measured with CellTiter 96 AQueous One Solution (MTS, Promega, #G3580). The assay was performed according to the methods described in the manufacturer’s manual. Briefly, cells (1 × 10^3^ cells) were seeded in 96-well plates and incubated in a standard incubator. At defined time points, 20 μL of CellTiter 96 AQueous One Solution Reagent was added and incubated for 2 hours at 37 °C. The quantity of formazan product, which is directly proportional to the number of living cells in the culture, was measured by absorbance at 490 nm with a 96-well plate reader. In the treatment of AOAA (CBS inhibitor, Sigma-Aldrich), cells (5 × 10^3^ cells) were seeded in 96-well plates overnight, treated with AOAA (0, 0.25, or 0.5 mM), and incubated in a standard incubator. At defined time points, cells were fixed in 4% formaldehyde for 30 min and permeabilized with 0.1% Triton X-100 in PBS for 10 min. 1 μg/mL DAPI (diluted in PBS, Sigma-Aldrich) was used to stain cells for cell counting.

### Glycerol gradient ultracentrifugation

To perform glycerol gradient ultracentrifugation, 150 μg cell lysates or 10 μg recombinant PKM2 proteins were loaded on top of 15-35% glycerol gradient. Following centrifugation at 50,000 rpm for 16 hours at 4 °C in SW 55 Ti Rotor (Beckman Coulter), fractions (100 μL each) were taken from top to bottom and the oligomerization of PKM2 was analyzed by Western blot.

### PI staining

MDA-MB-231 stable cell lines (2 × 10^6^ cells) were fixed in 1 mL PBS and 9 mL 70% ethanol and stored at −30 °C overnight, while MDA-MB-231 cells were pre-incubated 48 or 72 hours with 0.25 mM AOAA or H_2_O control before fixation. Cells were collected through centrifugation, washed, and resuspended in 500 μL PI/Triton X-100 staining solution [0.1% (v/v) Triton X-100 (Sigma-Aldrich) in PBS add 0.1 mg DNAse-free RNAse A (Sigma-Aldrich) and 10 μg PI (Sigma-Aldrich)]. The stained samples were incubated for 30 min in the dark at room temperature. After PI staining, the stained samples were analyzed using CytoFLEX (Beckman Coulter). Data were collected in 50,000 particle units and used to form a histogram of fluorescence area signals and analyzed by FlowJo software.

### Senescence Associated β-galactosidase (SA-β-gal) staining

MDA-MB-231 stable cell lines (5 × 10^3^ cells) were seeded in a 24-well plate and incubated 4 days before staining. Cells were fixed with 4% formaldehyde and stained with SA-β-gal staining solution [0.1% X-gal, 5 mM potassium ferrocyanide, 5 mM potassium ferricyanide, 150 mM NaCl, and 2 mM MgCl_2_ (SHOWA chemical, #1301-7260) in 40 mM citric acid/sodium phosphate solution, pH 6.0] for 3 days. Bright-field images were captured with 100X magnification.

### Cell cycle synchronization

MDA-MB-231 cells were transfected with vector control, PKM2^wt^, or PKM2^C326S^ mutant 24 hours before cell cycle synchronization. Cells were synchronized to G2/M transition with double thymidine block (2 mM) and RO-3306 (2.5 μg/mL), and then released in culture medium for 6 hours to arrest in metaphase.

### Immunoprecipitation (IP)

Cells were lysed in non-denaturing lysis buffer [20 mM Tris-HCl pH 8, 137 mM NaCl, 1% NP-40, 2 mM EDTA, and 1X protease inhibitor (Roche)]. The cell lysates were then centrifuged, and the supernatants were collected and diluted to the same concentration. The cell lysates (500 μL) were mixed with 0.5 μg anti-V5-tag antibody (CST), anti-Bub3 antibody (Abcam) or IgG control (Millipore), and 50 μL protein A agarose bead slurry (25 μL packed beads) or Protein A Mag Sepharose Xtra (Cytiva). The mixture was gently rocked overnight at 4 °C on the rocker. After the co-IP, the beads were removed from each sample and mixed with 2X sample buffer for western blot analysis.

### *In vivo* xenograft mouse model

The female SCID mice (C.B17/Icr-*Prkdc^scid^*/CrlNarl, NARLabs) were randomly assigned to three groups after arriving and injected with 2 × 10^6^ luciferase stable expressed breast cancer cells (MDA-MB-231 with GFP-luc-2A plus pLX304 or pLX304-PKM2, or pLX304-PKM2-C326S) in 100 µl PBS containing Matrigel (Corning, #356237) per mouse at the 4^th^ mammary fat pad at age of 8 to 10 weeks. The cells’ luciferase activity (Promega, #E1483 or PerkinElmer, #6016716) had been confirmed before injection. Mice were measured for body weight and tumor size once per week. Once the largest diameter of the largest tumor reached 20 mm, mice were intraperitoneally (IP) injected with luciferin (Promega, #P1043) for *in vivo* imaging system (IVIS, Perkin Elmer). After that, mice were sacrificed, and tumors were harvested for IVIS. All procedures used in our mouse study were performed according to the approved protocol by the Institutional Animal Care and Use Committee of National Tsing Hua University (NTHU-IACUC-10656).

### Gel filtration analysis

200 μL of purified protein samples at 0.2 mg/mL were injected into a Superdex 200 10/300 GL column (Cytiva) equilibrated in PBS buffer at 4 °C. Protein profiles were monitored by the absorbance at 280 nm.

### Crystallization and data collection

For crystallization of the PKM2^C326S^, purified protein was concentrated to 8 mg/mL in 25 mM Tris (pH 8.0, APOLO Biochemical, #APL-0082), 100 mM KCl (Sigma-Aldrich, #P5405), 1 mM DL-Dithiothreitol (DTT, UniRegion Bio-Tech, #UR-DTT), 10% glycerol (J.T.Baker, #2136-08). Protein crystals were grown in 0.4 M ammonium phosphate monobasic, 23% polyethylene glycerol 3350, Tris pH 7.0 at 22 °C by hanging drop vapor diffusion method and further improved by microseeding. The diffraction data were collected at beamline BL07A of the National Synchrotron Radiation Research Center (Taiwan), merged from several crystals, and processed by HKL2000 ^52^.

### Structural determination and refinement

The crystal structure of PKM2^C326S^ was solved by molecular replacement in Phenix ^53–55^ using wild-type PKM2 (PDB ID: 4QG6) without the B domain as the search model. Structural refinement and model building were carried out by Phenix ^53–55^ and Coot ^56^ iteratively. Structural analysis and presentation were performed by PyMOL ^57^.

### Clinical analysis from The Cancer Genome Atlas

For gene expression analysis, data containing 1033 breast cancer (BRCA) cases and 59 normal breast tissues were collected from The Cancer Genome Atlas (TCGA) database (https://portal.gdc.cancer.gov/projects/TCGA-BRCA). Gene expressions were represented as transcripts per million (TPM). After data mining, the student’s t-test was performed in R (The R Foundation, v 4.2.1). For Pearson correlation analysis, data collected from TCGA-BRCA were applied, and gene expressions were converted to log2(TPM). After data mining, Pearson correlation analyses were performed in R (v 4.2.1). For survival, Kaplan-Meier Plotter (http://kmplot.com/analysis/) was used to assess the prognostic value of CBS from TCGA-BRCA. The hazard ratios (HRs) with 95% confidence intervals (CIs) and log-rank p-values were also computed.

### Quantification and statistical analysis

The statistical details of the experiments are described in figure legends. Most statistical analyses were performed using GraphPad Prism software (GraphPad Prism, CA, USA) with student’s t-test to compare two means. ANOVA followed by Tukey’s post hoc test was used for the statistical analysis when more than two means were compared (*p< 0.05; **p< 0.01; ***p< 0.001; ****p< 0.0001).

## Acknowledgments

Mass spectrometry analyses were performed by the Research Facility for Sharing for Proteomics and Chemistry located at the Institute of Biological Chemistry, NHRI. LTQ-Orbitrap data were acquired at the Academia Sinica Common Mass Spectrometry Facilities for Proteomics and Protein Modification Analysis located at the Institute of Biological Chemistry, Academia Sinica, supported by Academia Sinica Core Facility and Innovative Instrument Project (AS-CFII-111-209). We thank the experimental facilities and the technical services provided by the “Synchrotron Radiation Protein Crystallography Facility of the National Core Facility Program for Biotechnology, NSTC”, the “National Synchrotron Radiation Research Center”, a national user facility supported by the NSTC, and the “Macromolecular Crystallography Center” at NTHU. We thank the technical support at the confocal imaging core at National Tsing Hua University.

## Data availability

The coordinates and structure factors of the crystal structure of PKM2 C326S have been deposited in the Protein Data Bank under PDB ID: 8HGF. The proteomic data have been deposited in the PRoteomics IDEntifications (PRIDE) database (Project accession: PXD044925).

## Materials availability

This study did not generate new unique reagents.

## Author contributions

Conceptualization: L-HW, H-CC, K-TL

Data curation: R-HW, P-RC, Y-TC, Y-CC, Y-HC, C-CC, P-CC, S-YL, Z-LW, M-CT, S-YC, G-SC, W-LC, Y-HW, S-YL, H-CC, K-TL

Resources: LW, W-CW, H-JK

Investigation: L-HW, H-CC, K-TL

Funding acquisition: H-CC, K-TL

Project administration: L-HW, H-CC, K-TL

Supervision: L-HW, H-CC, K-TL

Writing – original draft: R-HW, Y-CC, Y-HC, H-CC, K-TL

Writing – review & editing: LW, W-CW, H-JK, L-HW, H-CC, K-TL

## Ethics declarations

Competing interests: The authors declare no competing interests.

## References

1 Cairns, R. A., Harris, I. S. & Mak, T. W. Regulation of cancer cell metabolism. Nat Rev Cancer 11, 85–95 (2011). 10.1038/nrc2981

2 Warburg, O., Wind, F. & Negelein, E. The Metabolism of Tumors in the Body. J Gen Physiol 8, 519–530 (1927).

3 Pavlova, N. N. & Thompson, C. B. The Emerging Hallmarks of Cancer Metabolism. Cell Metab 23, 27–47 (2016). 10.1016/j.cmet.2015.12.006

4 Le, A. et al. Inhibition of lactate dehydrogenase A induces oxidative stress and inhibits tumor progression. Proc Natl Acad Sci U S A 107, 2037–2042 (2010). 10.1073/pnas.0914433107

5 Jones, R. S. & Morris, M. E. Monocarboxylate Transporters: Therapeutic Targets and Prognostic Factors in Disease. Clin Pharmacol Ther 100, 454–463 (2016). 10.1002/cpt.418

6 Gupta, V. & Bamezai, R. N. Human pyruvate kinase M2: a multifunctional protein. Protein Sci 19, 2031–2044 (2010). 10.1002/pro.505

7 Mazurek, S. Pyruvate kinase type M2: a key regulator of the metabolic budget system in tumor cells. Int J Biochem Cell Biol 43, 969–980 (2011). 10.1016/j.biocel.2010.02.005

8 Zhang, Z. et al. PKM2, function and expression and regulation. Cell Biosci 9, 52 (2019). 10.1186/s13578-019-0317-8

9 Yang, W. et al. ERK1/2-dependent phosphorylation and nuclear translocation of PKM2 promotes the Warburg effect. Nat Cell Biol 14, 1295–1304 (2012). 10.1038/ncb2629

10 Lv, L. et al. Acetylation targets the M2 isoform of pyruvate kinase for degradation through chaperone-mediated autophagy and promotes tumor growth. Mol Cell 42, 719–730 (2011). 10.1016/j.molcel.2011.04.025

11 Anastasiou, D. et al. Pyruvate kinase M2 activators promote tetramer formation and suppress tumorigenesis. Nat Chem Biol 8, 839–847 (2012). 10.1038/nchembio.1060

12 Spoden, G. A. et al. The SUMO-E3 ligase PIAS3 targets pyruvate kinase M2. J Cell Biochem 107, 293–302 (2009). 10.1002/jcb.22125

13 Wang, Y. et al. O-GlcNAcylation destabilizes the active tetrameric PKM2 to promote the Warburg effect. Proc Natl Acad Sci U S A 114, 13732–13737 (2017). 10.1073/pnas.1704145115

14 Liu, F. et al. PKM2 methylation by CARM1 activates aerobic glycolysis to promote tumorigenesis. Nat Cell Biol 19, 1358–1370 (2017). 10.1038/ncb3630

15 Amin, S., Yang, P. & Li, Z. Pyruvate kinase M2: A multifarious enzyme in non-canonical localization to promote cancer progression. Biochim Biophys Acta Rev Cancer 1871, 331–341 (2019). 10.1016/j.bbcan.2019.02.003

16 Szabo, C. Hydrogen sulphide and its therapeutic potential. Nat Rev Drug Discov 6, 917–935 (2007). 10.1038/nrd2425

17 Coletta, C. et al. Hydrogen sulfide and nitric oxide are mutually dependent in the regulation of angiogenesis and endothelium-dependent vasorelaxation. Proc Natl Acad Sci U S A 109, 9161–9166 (2012). 10.1073/pnas.1202916109

18 Mustafa, A. K. et al. Hydrogen sulfide as endothelium-derived hyperpolarizing factor sulfhydrates potassium channels. Circ Res 109, 1259–1268 (2011). 10.1161/CIRCRESAHA.111.240242

19 Yang, G. et al. H2S as a physiologic vasorelaxant: hypertension in mice with deletion of cystathionine gamma-lyase. Science 322, 587–590 (2008). 10.1126/science.1162667

20 Hine, C. et al. Endogenous hydrogen sulfide production is essential for dietary restriction benefits. Cell 160, 132–144 (2015). 10.1016/j.cell.2014.11.048

21 Zanardo, R. C. et al. Hydrogen sulfide is an endogenous modulator of leukocyte-mediated inflammation. FASEB J 20, 2118–2120 (2006). 10.1096/fj.06-6270fje

22 Wang, R. H., Chu, Y. H. & Lin, K. T. The Hidden Role of Hydrogen Sulfide Metabolism in Cancer. Int J Mol Sci 22 (2021). 10.3390/ijms22126562

23 Paul, B. D. & Snyder, S. H. H(2)S signalling through protein sulfhydration and beyond. Nat Rev Mol Cell Biol 13, 499–507 (2012). 10.1038/nrm3391

24 Nakatsu, D. et al. L-cysteine reversibly inhibits glucose-induced biphasic insulin secretion and ATP production by inactivating PKM2. Proc Natl Acad Sci U S A 112, E1067–1076 (2015). 10.1073/pnas.1417197112

25 Kimura, H. Hydrogen sulfide and polysulfides as signaling molecules. Proc Jpn Acad Ser B Phys Biol Sci 91, 131–159 (2015). 10.2183/pjab.91.131

26 Luo, W. & Semenza, G. L. Emerging roles of PKM2 in cell metabolism and cancer progression. Trends Endocrinol Metab 23, 560–566 (2012). 10.1016/j.tem.2012.06.010

27 Xiao, H. et al. PKM2 Promotes Breast Cancer Progression by Regulating Epithelial Mesenchymal Transition. Anal Cell Pathol (Amst) 2020, 8396023 (2020). 10.1155/2020/8396023

28 Anastasiou, D. et al. Inhibition of pyruvate kinase M2 by reactive oxygen species contributes to cellular antioxidant responses. Science 334, 1278–1283 (2011). 10.1126/science.1211485

29 Zivanovic, J. et al. Selective Persulfide Detection Reveals Evolutionarily Conserved Antiaging Effects of S-Sulfhydration. Cell Metab 30, 1152–1170 e1113 (2019). 10.1016/j.cmet.2019.10.007

30 Paulsen, C. E. & Carroll, K. S. Cysteine-mediated redox signaling: chemistry, biology, and tools for discovery. Chem Rev 113, 4633–4679 (2013). 10.1021/cr300163e

31 Morgan, H. P. et al. M2 pyruvate kinase provides a mechanism for nutrient sensing and regulation of cell proliferation. Proc Natl Acad Sci U S A 110, 5881–5886 (2013). 10.1073/pnas.1217157110

32 Wang, P., Sun, C., Zhu, T. & Xu, Y. Structural insight into mechanisms for dynamic regulation of PKM2. Protein Cell 6, 275–287 (2015). 10.1007/s13238-015-0132-x

33 Chaneton, B. et al. Serine is a natural ligand and allosteric activator of pyruvate kinase M2. Nature 491, 458–462 (2012). 10.1038/nature11540

34 Icard, P., Fournel, L., Wu, Z., Alifano, M. & Lincet, H. Interconnection between Metabolism and Cell Cycle in Cancer. Trends Biochem Sci 44, 490–501 (2019). 10.1016/j.tibs.2018.12.007

35 Yang, W. et al. PKM2 phosphorylates histone H3 and promotes gene transcription and tumorigenesis. Cell 150, 685–696 (2012). 10.1016/j.cell.2012.07.018

36 Jiang, Y. et al. PKM2 regulates chromosome segregation and mitosis progression of tumor cells. Mol Cell 53, 75–87 (2014). 10.1016/j.molcel.2013.11.001

37 Kuilman, T., Michaloglou, C., Mooi, W. J. & Peeper, D. S. The essence of senescence. Genes Dev 24, 2463–2479 (2010). 10.1101/gad.1971610

38 Jiang, Y. et al. PKM2 phosphorylates MLC2 and regulates cytokinesis of tumour cells. Nat Commun 5, 5566 (2014). 10.1038/ncomms6566

39 Cuevasanta, E. et al. Reaction of Hydrogen Sulfide with Disulfide and Sulfenic Acid to Form the Strongly Nucleophilic Persulfide. J Biol Chem 290, 26866–26880 (2015). 10.1074/jbc.M115.672816

40 Boxer, M. B. et al. Evaluation of substituted N,N’-diarylsulfonamides as activators of the tumor cell specific M2 isoform of pyruvate kinase. J Med Chem 53, 1048–1055 (2010). 10.1021/jm901577g

41 Jiang, J. K. et al. Evaluation of thieno[3,2-b]pyrrole[3,2-d]pyridazinones as activators of the tumor cell specific M2 isoform of pyruvate kinase. Bioorg Med Chem Lett 20, 3387–3393 (2010). 10.1016/j.bmcl.2010.04.015

42 Barnum, K. J. & O’Connell, M. J. Cell cycle regulation by checkpoints. Methods Mol Biol 1170, 29–40 (2014). 10.1007/978-1-4939-0888-2_2

43 Musacchio, A. & Salmon, E. D. The spindle-assembly checkpoint in space and time. Nat Rev Mol Cell Biol 8, 379–393 (2007). 10.1038/nrm2163

44 Holland, A. J. & Cleveland, D. W. Boveri revisited: chromosomal instability, aneuploidy and tumorigenesis. Nat Rev Mol Cell Biol 10, 478–487 (2009). 10.1038/nrm2718

45 Shin, E. & Koo, J. S. Glucose Metabolism and Glucose Transporters in Breast Cancer. Front Cell Dev Biol 9, 728759 (2021). 10.3389/fcell.2021.728759

46 Tseng, C. W. et al. Transketolase Regulates the Metabolic Switch to Control Breast Cancer Cell Metastasis via the alpha-Ketoglutarate Signaling Pathway. Cancer Res 78, 2799–2812 (2018). 10.1158/0008-5472.CAN-17-2906

47 Woo, Y. M. et al. Inhibition of Aerobic Glycolysis Represses Akt/mTOR/HIF-1alpha Axis and Restores Tamoxifen Sensitivity in Antiestrogen-Resistant Breast Cancer Cells. PLoS One 10, e0132285 (2015). 10.1371/journal.pone.0132285

48 Zhao, Y. et al. Overcoming trastuzumab resistance in breast cancer by targeting dysregulated glucose metabolism. Cancer Res 71, 4585–4597 (2011). 10.1158/0008-5472.CAN-11-0127

49 Longchamp, A. et al. Amino Acid Restriction Triggers Angiogenesis via GCN2/ATF4 Regulation of VEGF and H(2)S Production. Cell 173, 117–129 e114 (2018). 10.1016/j.cell.2018.03.001

50 Schneider, C. A., Rasband, W. S. & Eliceiri, K. W. NIH Image to ImageJ: 25 years of image analysis. Nat Methods 9, 671–675 (2012). 10.1038/nmeth.2089

51 Mustafa, A. K. et al. H2S signals through protein S-sulfhydration. Sci Signal 2, ra72 (2009). 10.1126/scisignal.2000464

52 Otwinowski, Z. & Minor, W. Processing of X-ray diffraction data collected in oscillation mode. Methods Enzymol 276, 307–326 (1997). 10.1016/S0076-6879(97)76066-X

53 Adams, P. D. et al. PHENIX: building new software for automated crystallographic structure determination. Acta Crystallogr D Biol Crystallogr 58, 1948–1954 (2002). 10.1107/s0907444902016657

54 Adams, P. D. et al. PHENIX: a comprehensive Python-based system for macromolecular structure solution. Acta Crystallogr D Biol Crystallogr 66, 213–221 (2010). 10.1107/S0907444909052925

55 Afonine, P. V. et al. Towards automated crystallographic structure refinement with phenix.refine. Acta Crystallogr D Biol Crystallogr 68, 352–367 (2012). 10.1107/S0907444912001308

56 Emsley, P. & Cowtan, K. Coot: model-building tools for molecular graphics. Acta Crystallogr D Biol Crystallogr 60, 2126–2132 (2004). 10.1107/S0907444904019158

57 Schrodinger, LLC. The PyMOL Molecular Graphics System, Version 18. (2015).

