## Supplemental Information for "Hydrogen Sulfide Coordinates Glucose Metabolism Switch through Destabilizing Tetrameric Pyruvate Kinase M2"

**The file includes**

Table S1-S3

Figure S1-S6

**Other Supplementary Material for this manuscript includes the following:**

**Movie S1 -** Time-lapse movie of cell division in MDA-MB-231 cells expressing wildtype PKM2.

**Movie S2 -** Time-lapse movie of cell division in MDA-MB-231 cells expressing $PKM2^{C326S}$.

**Table S1. Overview of cysteine PTMs on PKM2 detected by LC-MS/MS.**

| **Modification** | **Formulas** | **Monoisotopic mass change** | **Detectable** | **Sites** |
|---|---|---|---|---|
| Beta-methylthiolation | S-SCH3<br>(SH modified by MMTS) | + 45.988 | V | Cys 31, 49, 152, 326, 358, 423, 424, and 474 |
| SSCAM | S-C2H3NOS<br>(SSH modified by IAM) | + 88.994 | V | Cys 326 |
| Carbamidomethylation (CAM) | S-C2H3NO<br>(SH modified by IAM) | + 57.021 | X | |
| S-Nitrosylation | S-NO | +28.990 | X | |
| Glutathionylation | S-SG | +305.068 | X | |
| Oxidation | S-OH | +15.995 | X | |
| Dioxidation* | S-SO2H | +31.990 | X | |
| Trioxidation* | S-SO3H | +47.985 | X | |

Analysis of PTMs identified from cell lysates of MDA-MB-231. Related to Figure 2B.
*Irreversible Modifications

**Table S2. Data collection and refinement statistics.**

| | PKM2 C326S |
|---|---|
| **Data collection** | |
| Wavelength (Å) | 0.97626 |
| Resolution range (Å) | 29.69 - 3.10 (3.21 - 3.10) |
| Space group | C 1 2 1 |
| Unit cell a, b, c (Å); α, β, γ (°) | 181.51, 156.37, 121.59; 90, 114.20, 90 |
| Total reflections | 55273 (5394) |
| Unique reflections | 15354 (1498) |
| Multiplicity | 3.6 (3.6) |
| Completeness (%) | 98.4 (96.5) |
| Mean I/sigma (I) | 20.0 (2.4) |
| R-meas (%) | 6.8 (60.1) |
| CC1/2 | 0.998 (0.877) |
| **Refinement** | |
| Reflections used in refinement | 49956 (3326) |
| Reflections used for R-free | 1999 (133) |
| R-work (%) | 18.4 (29.3) |
| R-free (%) | 23.8 (34.9) |
| Number of non-hydrogen atoms | |
| macromolecules | 14857 |
| Protein residues | 1938 |
| RMS(bonds) (Å) | 0.022 |
| RMS(angles) (°) | 1.86 |
| Average B-factor (Å2) | 55.31 |

Statistics for the highest-resolution shell are shown in parentheses.

**Table S3. Lists of primers used in this study.**

.

| | |
|---|---|
| CCND1 | Forward: GCGAGGAACAGAAGTG |
| | Reverse: GAGTTGTCGGTGTAGATGC |
| HIF-1α | Forward: GCCGAGGAAGAACTATGA |
| | Reverse: GGTTGGTTACTGTTGGTATC |
| LDHA | Forward: CCAGAATAAGATTACAGTTGTTG |
| | Reverse: GCCAAGTCCTTCATTAAGATA |
| GLUT1 | Forward: GCAGGAGATGAAGGAAGA |
| | Reverse: AATAGAAGACAGCGTTGATG |
| GLUT12 | Forward: TTGACTGTAACTGATCTTATTGG |
| | Reverse: ACTAATTCTTCTTGGTGATGAC |
| HK2 | Forward: TTGACCAGGAGATTGACAT |
| | Reverse: GCCATCTTCACCAGGATA |
| GLS1 | Forward: TTTGTGATTCCTGACTTTATGT |
| | Reverse: GCCTCTGTCCATCTACTG |

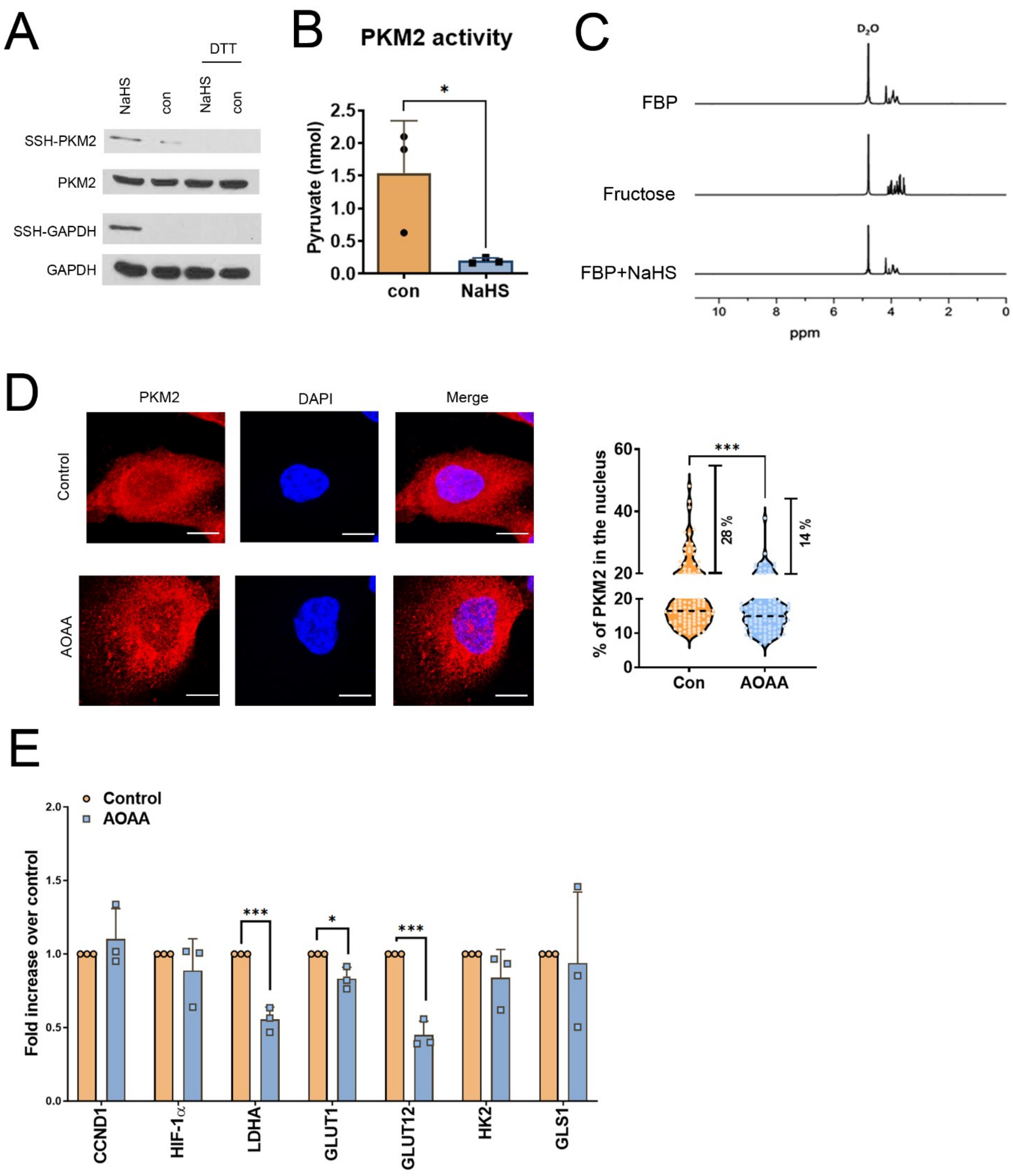
A
DTT
NaHS
con
NaHS
con
SSH-PKM2
PKM2
SSH-GAPDH
GAPDH
B
PKM2 activity
Pyruvate (nmol)
con
NaHS
C
D₂O
FBP
Fructose
FBP+NaHS
ppm
D
PKM2
DAPI
Merge
Control
AOAA
% of PKM2 in the nucleus
28 %
14 %
Con
AOAA
E
Control
AOAA
Fold increase over control
CCND1
HIF-1α
LDHA
GLUT1
GLUT12
HK2
GLS1

**Figure S1. $H_2S$ mediates PKM2 sulfhydration to inhibit PK activity.** (A) PC3 cell lysates were treated with 100 μM NaHS for 30 min at 37°C and subjected to 1 mM DTT for 10 min. The biotin switch assay was then applied to precipitate sulfhydrated proteins. The biotin-labeled protein was analyzed by immunoblotting with anti-PKM2 or GAPDH antibody to detect sulfhydration of PKM2 and GAPDH (as positive control). (B) PC3 cell lysates were treated with 1 μM NaHS for 30 min at 37°C. Cells were lysed and pyruvate kinase activities were then assayed by measuring the amount of pyruvate production (nmol) (n=3 biological replicates). The Student's t-test was used for the statistical analysis (*$p<0.05$). (C) $^1$H NMR spectra of 10 mg FBP, 10 mg Fructose, or 10 mg FBP co-treated with 1 mg NaHS. (D) Left: MDA-MB-231 cells were exposed to 0.25 mM AOAA for 24h. Subcellular localization of PKM2 was detected by immunocytochemistry. Nuclei were counterstained with DAPI. The representative images are shown. Scale bars: 10 μm. Right: The percentage of nuclear PKM2 is shown in a violin plot with individual points. Horizontal black dotted lines display the median and the percentage of cells with >20% nuclear localization of PKM2 were indicated (n= 95-97 individual cells, data were combined from three independent experiments). The student’s t-test was used for the statistical analysis (***$p<0.001$). (E) MDA-MB-231 cells were treated with 0.25 mM AOAA for 24h. The relative expression of each gene was measured by qRT-PCR. Data are presented as the means ± SD (n=3 biological replicates). The student’s t-test was used for the statistical analysis (*$p<0.05$; ***$p<0.001$).

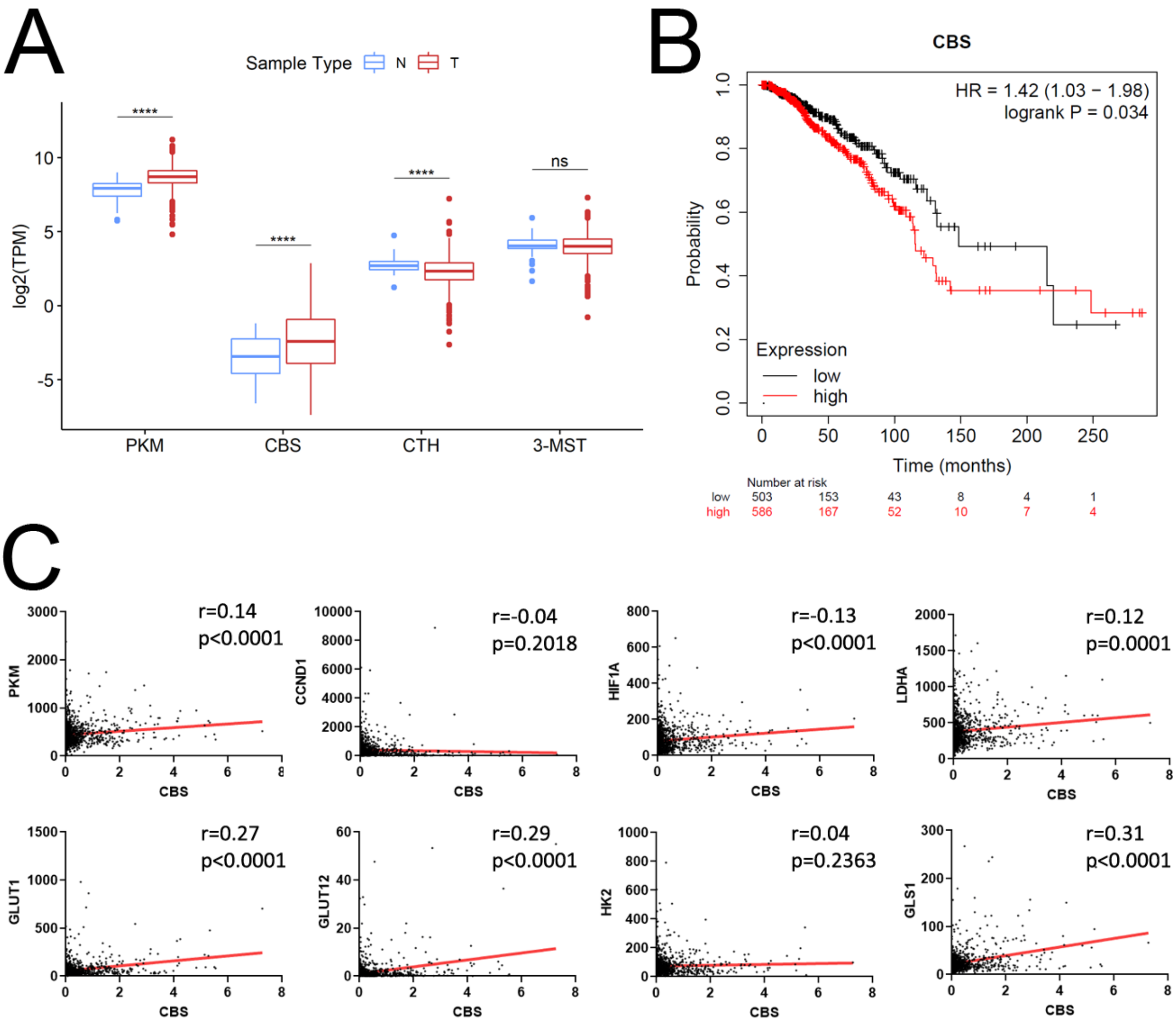
A
Sample Type N T
log2(TPM)
****
****
****
ns
PKM
CBS
CTH
3-MST
B
CBS
HR = 1.42 (1.03 − 1.98)
logrank P = 0.034
Probability
Expression
low
high
Time (months)
Number at risk
low 503 153 43 8 4 1
high 586 167 52 10 7 4
C
r=0.14
p<0.0001
PKM
r=-0.04
p=0.2018
CCND1
r=-0.13
p<0.0001
HIF1A
r=0.12
p=0.0001
LDHA
r=0.27
p<0.0001
GLUT1
r=0.29
p<0.0001
GLUT12
r=0.04
p=0.2363
HK2
r=0.31
p<0.0001
GLS1
CBS

**Figure S2. The expression, prognosis, and correlation of PKM and $H_2S$ synthesizing enzymes in breast cancer patients.** (A) Box plots represent gene expressions of PKM, CBS, CTH, and 3-MST between the normal breast tissues (N; n = 60) and breast tumors (T; n=1034) from the TCGA-BRCA RNA-seq dataset, measured in TPMs. The student's t-test was used for the statistical analysis (****$p$<0.0001; ns, not significant). (B) Kaplan-Meier overall survival analysis of breast cancer patients presenting high (n=586) or low (n=503) CBS expression. The RNA-seq dataset was from TCGA cohorts. The p-value was calculated using the log-rank test. (C) Plot of Pearson correlation of the expression level for CBS versus PKM or PKM2 responsive genes, including CCND1, HIF1A, LDHA, GLUT1, GLUT12, HK2, and GLS1.

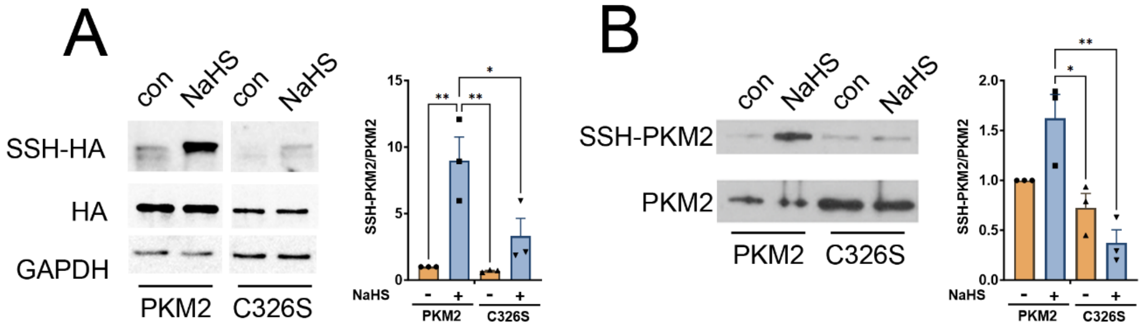

A
con
NaHS
con
NaHS
SSH-HA
HA
GAPDH
PKM2
C326S
SSH-PKM2/PKM2
15
10
5
0
**
**
*
NaHS
-
+
-
+
PKM2
C326S
B
con
NaHS
con
NaHS
SSH-PKM2
PKM2
PKM2
C326S
SSH-PKM2/PKM2
2.0
1.5
1.0
0.5
0.0
**
*
NaHS
-
+
-
+
PKM2
C326S

**Figure S3. PKM2 is sulfhydrated by $H_2S$, notably cysteine 326.** (A) Left: The cell lysate from MDA-MB-231 cells transfected with HA-tagged PKM2 or $PKM2^{C326S}$ mutants was treated with 100 μM NaHS for 30 min at 37°C. The modified biotin switch assay was then applied to precipitate sulfhydrated protein. The biotin-labeled protein was analyzed by immunoblotting with anti-HA antibodies to detect sulfhydration of transfected PKM2. Right: The SSH-labeled HA-PKM2 was normalized with the level of total HA-PKM2. Data are presented as the means ± SEM (n=3 biological replicates). One-way ANOVA followed by post hoc test was used for the statistical analysis (*p<0.05; **p<0.01). (B) Left: Purified recombinant PKM2 or $PKM2^{C326S}$ mutant protein was treated with 100 μM NaHS for 30 min at 37°C. The modified biotin switch assay was then applied to precipitate sulfhydrated protein. The biotin-labeled protein was analyzed by immunoblotting with anti-PKM2 antibody to detect sulfhydration of PKM2. Right: The SSH-labeled PKM2 was normalized with the level of total PKM2. Data are presented as the means ± SEM (n=3 biological replicates). One-way ANOVA followed by post hoc test was used for the statistical analysis (*p<0.05; **p<0.01).

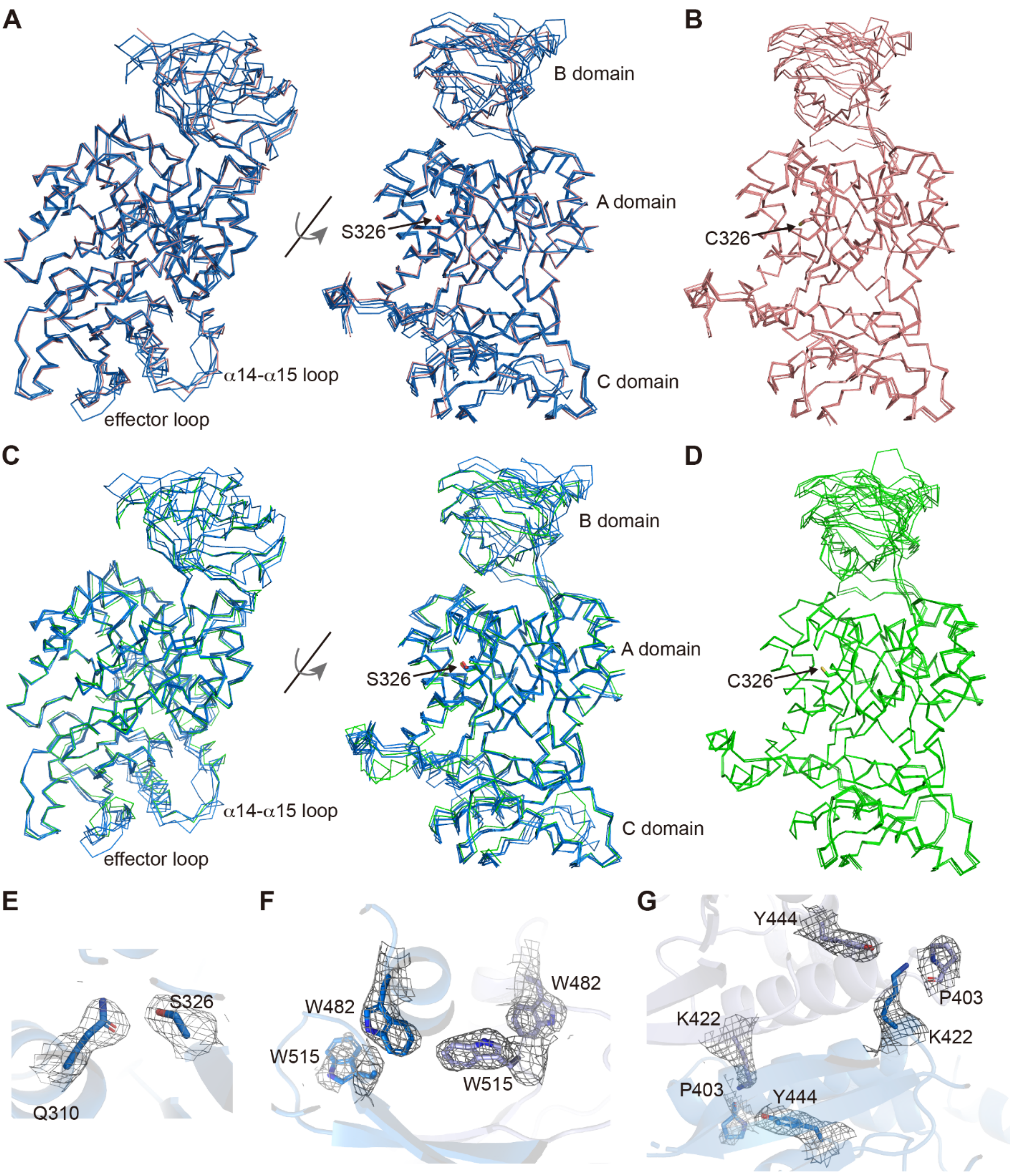

A
B domain
A domain
S326
C domain
α14-α15 loop
effector loop
B
C326
C
B domain
A domain
S326
C domain
α14-α15 loop
effector loop
D
C326
E
S326
Q310
F
W482
W515
W482
W515
G
Y444
P403
K422
K422
P403
Y444

**Figure S4. Structural analyses of PKM2$^{C326S,}$ compared to the T and R states.** (A) Structural overlay of PKM2$^{C326S}$ protomers on one of the T-state protomers. (B) Structural overlay of PKM2 T-state protomers (PDB: 4FXJ). (C) Structural overlay of PKM2$^{C326S}$ protomers on one of the R-state protomers. (D) Structural overlay of PKM2 R-state protomers (PDB: 4B2D). PKM2$^{C326S}$ protomers are in blue, T-state protomers in pink, and R-state protomers in green. (E-G) 2Fo-Fc electron density map contoured at 1 σ for the designated residues.

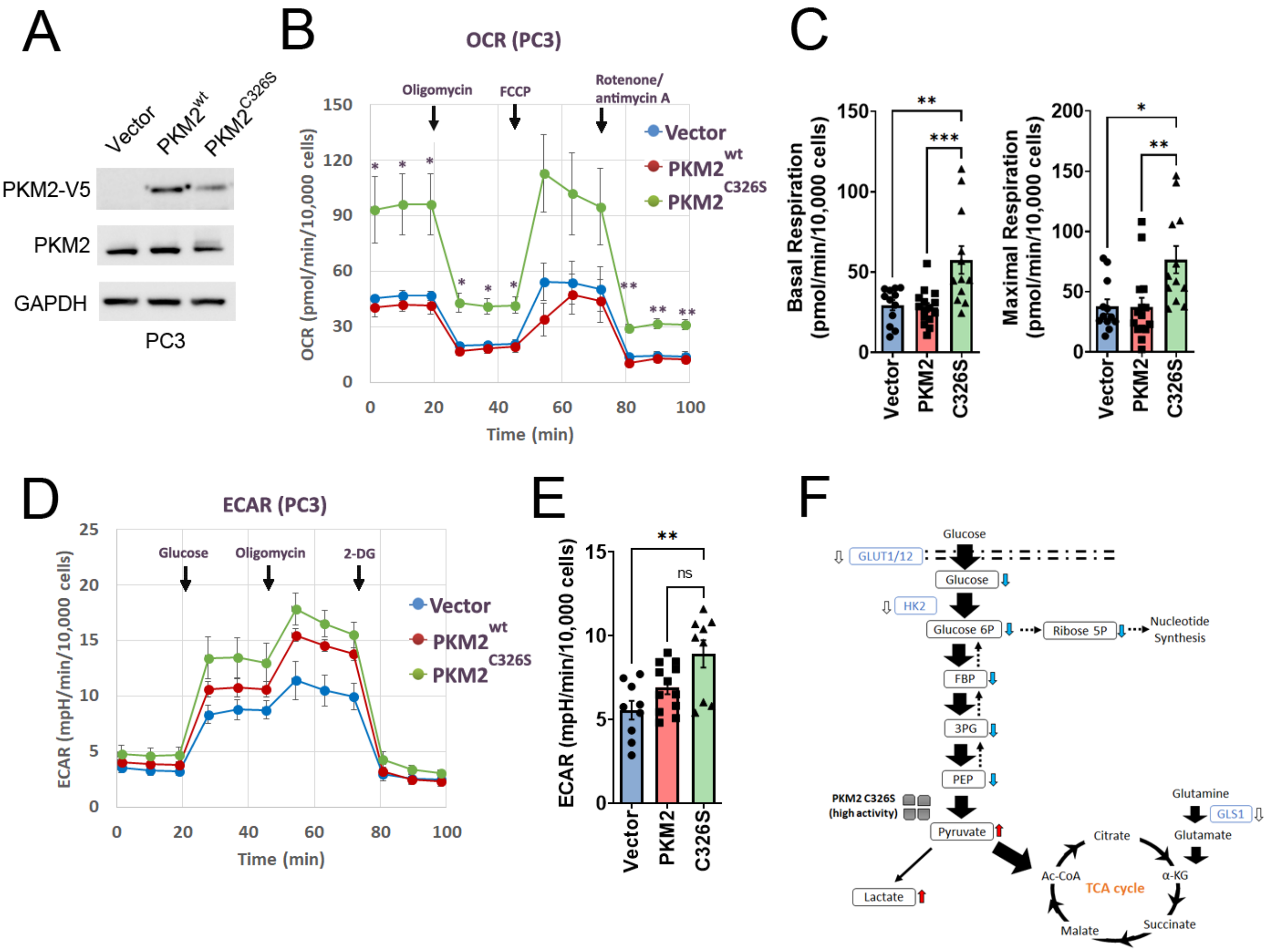
A
Vector
PKM2wt
PKM2C326S
PKM2-V5
PKM2
GAPDH
PC3
B
OCR (PC3)
Oligomycin
FCCP
Rotenone/
antimycin A
Vector
PKM2wt
PKM2C326S
OCR (pmol/min/10,000 cells)
Time (min)
C
Basal Respiration
(pmol/min/10,000 cells)
Maximal Respiration
(pmol/min/10,000 cells)
Vector
PKM2
C326S
ns
D
ECAR (PC3)
Glucose
Oligomycin
2-DG
Vector
PKM2wt
PKM2C326S
ECAR (mpH/min/10,000 cells)
Time (min)
E
ECAR (mpH/min/10,000 cells)
Vector
PKM2
C326S
F
Glucose
GLUT1/12
Glucose
HK2
Glucose 6P
Ribose 5P
Nucleotide
Synthesis
FBP
3PG
PEP
PKM2 C326S
(high activity)
Pyruvate
Lactate
Glutamine
GLS1
Glutamate
Citrate
Ac-CoA
α-KG
TCA cycle
Malate
Succinate

**Figure S5. Blockage of PKM2 C326 sulfhydration enhances mitochondrial oxidative phosphorylation.** (A) Western blot analysis of stable expression of V5-tagged PKM2 (wild type or C326S mutant) in PC3 cells. (B) The oxygen consumption rate (OCR) curves in PC3 cells expressing Vector alone, $PKM2^{wt}$, or $PKM2^{C326S}$. Cells were treated with oligomycin, FCCP, and rotenone/antimycin A, respectively (n=3 biological replicates). The student's t-test was used for the statistical analysis (**p*<0.05, ***p*<0.01; C326S compared to the vector or PKM2 group). (C) The level of basal OCR and maximal OCR normalized to the cell numbers in PC3 cells expressing Vector alone, $PKM2^{wt}$, or $PKM2^{C326S}$ (n=9 or 12 experiments per group, data were combined from three independent experiments). One-way ANOVA followed by post hoc test was used for the statistical analysis (**p*<0.05; ***p*<0.01; ****p*<0.001). (D) The ECAR curves in PC3 cells expressing Vector alone, $PKM2^{wt}$, or $PKM2^{C326S}$. Cells were treated with glucose, oligomycin, and 2-DG, respectively (n=3 biological replicates). (E) Glycolysis normalized to the cell numbers in PC3 cells expressing Vector alone, $PKM2^{wt}$, or $PKM2^{C326S}$ (n=9 or 12 experiments per group, data were combined from three independent experiments). One-way ANOVA followed by post hoc test was used for the statistical analysis (***p*<0.01; ns: not significant). (F) Schematic diagram of how blocking PKM2 sulfhydration at C326 facilitates the TCA cycle, resulting in the shortage of glycolytic intermediates for rapid DNA synthesis and enhanced oxidative phosphorylation.

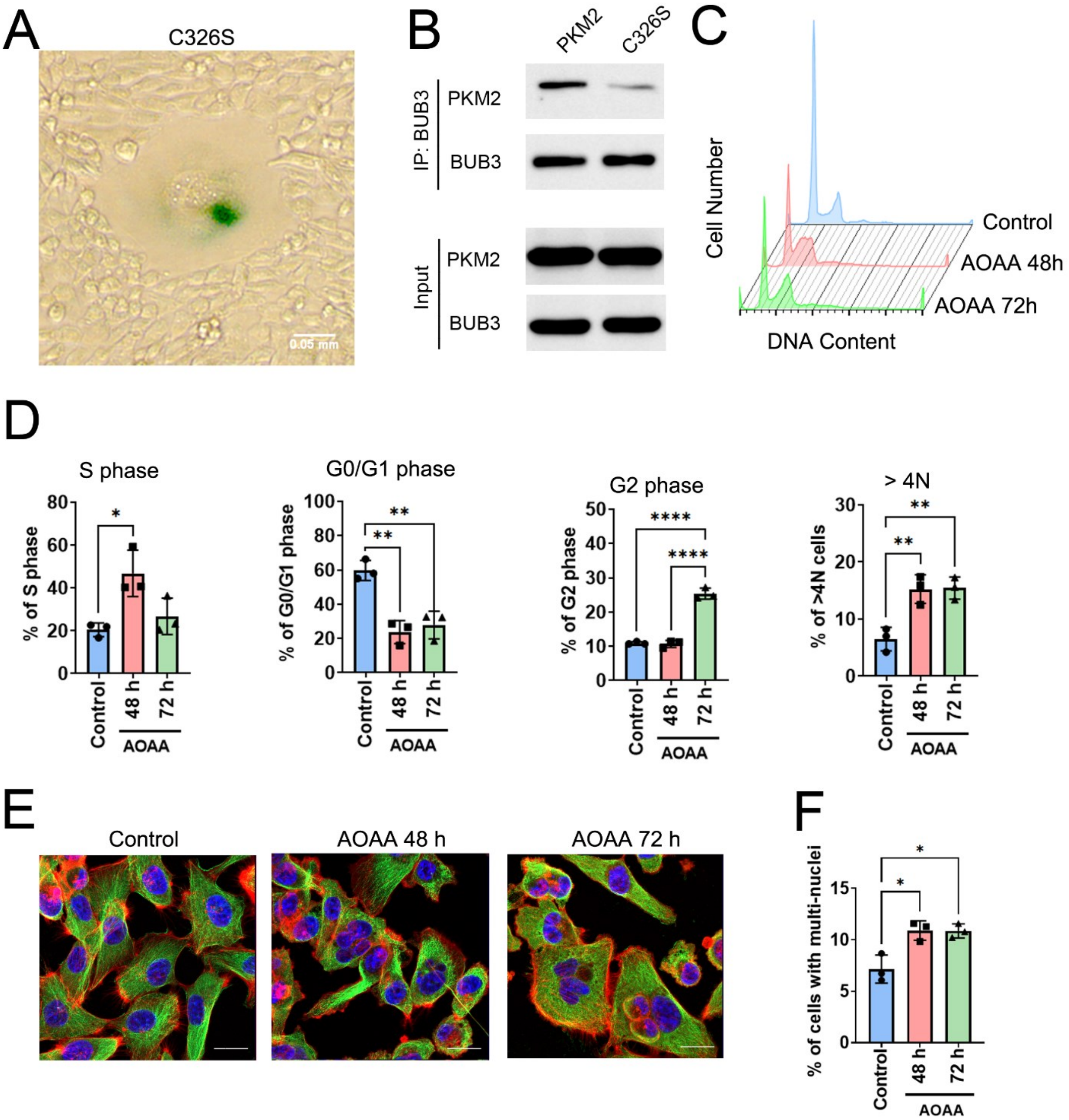
A
C326S
0.05 mm
B
PKM2
C326S
IP: BUB3
PKM2
BUB3
Input
PKM2
BUB3
C
Cell Number
Control
AOAA 48h
AOAA 72h
DNA Content
D
S phase
% of S phase
G0/G1 phase
% of G0/G1 phase
G2 phase
% of G2 phase
> 4N
% of >4N cells
Control
48 h
72 h
AOAA
E
Control
AOAA 48 h
AOAA 72 h
F
% of cells with multi-nuclei
Control
48 h
72 h
AOAA

**Figure S6. Blockage of PKM2 sulfhydration at C326 or depletion of $H_2S$ by AOAA causes defects in cell cycle progression and cytokinesis failure.** (A) SA-β-gal activity staining of MDA-MB-231 cells expressing $PKM2^{C326S}$. Scale bar: 0.05 mm. (B) MDA-MB-231 cells transfected with wildtype PKM2 or $PKM2^{C326S}$ were synchronized by double thymidine block (2 mM) and then with RO-3306 (2.5 μg/mL) for 12 hr. After being released for 6 h, cells entering metaphase were harvested, immunoprecipitated with anti-BUB3 antibody, and immunoblotted with PKM2 and BUB3 antibodies. (C, D) MDA-MB-231 cells were treated with 0.25 mM AOAA for 48 or 72 h. The cell cycle was examined by the PI staining method and measured by flow cytometry. (C) The representative image is shown. (D) The percentage of cells at different cell cycle phases was analyzed. Data are presented as the means ± SD (n=3 biological replicates). One-way ANOVA followed by post hoc test was used for the statistical analysis (*$p$<0.05; **$p$<0.01; ****$p$<0.0001). (E, F) Immunostaining of actin filaments (Red), α-tubulin (Green), and DNA counterstaining with DAPI (Blue) of MDA-MB-231 cells with 0.25 mM AOAA treatment for 48 or 72 h. (E) Representative images are shown. Scale Bar: 20μm. (F) The percentage of poly-nuclei cells (n≧2) was analyzed. Data are presented as the means ± SD (n=3 biological replicates). One-way ANOVA followed by post hoc test was used for the statistical analysis (*$p$<0.05).
